# *occAssess*: An R package for assessing potential biases in species occurrence data

**DOI:** 10.1101/2021.04.19.440441

**Authors:** R. J. Boyd, G. Powney, C. Carvell, O. L. Pescott

## Abstract

Species occurrence records from a variety of sources are increasingly aggregated into heterogeneous databases and made available to ecologists for immediate analytical use. However, these data are typically biased, i.e. they are not a probability sample of the target population of interest, meaning that the information they provide may not be an accurate reflection of reality. It is therefore crucial that species occurrence data are properly scrutinised before they are used for research. In this article, we introduce occAssess, an R package that enables straightforward screening of species occurrence data for potential biases. The package contains a number of discrete functions, each of which returns a measure of the potential for bias in one or more of the taxonomic, temporal, spatial and environmental dimensions. Users can opt to provide a set of time periods into which the data will be split; in this case separate outputs will be provided for each period, making the package particularly useful for assessing the suitability of a dataset for estimating temporal trends in species’ distributions. The outputs are provided visually (as ggplot2 objects) and do not include a formal recommendation as to whether data are of sufficient quality for any given inferential use. Instead, they should be used as ancillary information and viewed in the context of the question that is being asked, and the methods that are being used to answer it. We demonstrate the utility of occAssess by applying it to data on two key pollinator taxa in South America: leaf-nosed bats (Phyllostomidae) and hoverflies (Syrphidae). In this worked example, we briefly assess the degree to which various aspects of data coverage appear to have changed over time. We then discuss additional applications of the package, highlight its limitations, and point to future development opportunities.

## Introduction

Species occurrence records comprise information in three basic dimensions: taxonomic, geographic, and temporal; that is to say, what was seen, where was it seen, and when. Humans have been accumulating species occurrence data for centuries: historically as preserved specimens in museums and herbaria (Newbold, 2010; Spear et al., 2017), and in written accounts (e.g. Oswald and Preston, 2011); and more recently through recording for distribution atlases (Preston, 2013), and various other structured and unstructured monitoring and citizen science initiatives (Boakes et al., 2010; Pescott et al., 2015; Petersen et al., 2021). Taken together, these data provide an immense resource documenting species’ geographical distributions and opportunities to investigate how they may have changed over time. Over the last two decades, species occurrence data have become increasingly accessible, thanks to the digitisation of historic records and the launch of online data portals such as the Global Biodiversity Information Facility [GBIF; Nelson and Ellis (2019)]. A corollary of this increase in accessibility has been a surge in the use of species occurrence data for research in biodiversity conservation and other fields (Ball-Damerow et al., 2019).

Whilst clearly an increasingly important resource for ecologists, species occurrence data should be used with caution when drawing inferences about species’ distributions and how they have changed over time. Straightforward inference in statistics is predicated on the assumption that the data have been sampled randomly from the population of interest [probability sampling; e.g., Krzanowski (2010)]. In many, if not most, cases, species occurrence data available through aggregated databases do not satisfy this assumption. For example, data collected through citizen science initiatives tend to be collected opportunistically (sometimes called convenience sampling); that is, without a structured sampling plan. In this case, recorders are free to decide what to record, where and when. This generally leads to preferential sampling of attractive and accessible locations, and to documentation of interesting (e.g., rare) species (Isaac and Pocock, 2015). These “sampling biases” give rise to nonprobability samples which are not representative of the spatial, temporal and taxonomic domains of interest. Structured monitoring data tend to more closely resemble probability samples (although issues like sample dropout and patchy uptake may still create issues). However, when multiple structured datasets, with different aims, extents and sampling protocols, are aggregated (e.g. as on GBIF), the ultimate target population sampled by these activities is unlikely to be formally identified for inferential purposes. It may be possible to mitigate for biases by modifying the data [e.g., spatial thinning; Beck et al. (2014)] or through the use of statistical correction procedures (e.g., by modelling the data generation process; Turner et al., 2009). In order to decide on what mitigating action might be required, or if the data are simply too unrepresentative for a given use, it would be helpful to have a set of heuristics that can indicate the degree to which a dataset might suffer from various forms of bias.

There is a growing literature of studies which take species occurrence datasets and screen them for biases (Barends et al., 2020; Boakes et al., 2010; Meyer et al., 2016; Pescott et al., 2019a; Petersen et al., 2021; Ruete, 2015; Speed et al., 2018; Sumner et al., 2019; Troudet et al., 2018); we also note that various approaches to visualising the spatial and temporal coverage of occurrence records across large areas have been commonplace in national species atlases for some time (e.g. Preston et al., 2002). Studies of these types provide a template for how to conduct such assessments, and a suite of heuristics which can be deployed in similar situations. For example, one could assess data for spatial bias by comparing the nearest neighbour distances of the occurrence data with those from a simulated random distribution (Sumner et al., 2019). The proportion of records identified to species level can be used as a measure of how taxonomic uncertainty has changed over time (Troudet et al., 2018). Multidimensional environmental space can be summarised using principal component analyses (PCAs), or other ordination techniques, allowing one to map the distribution of records in environmental space and scrutinise it for bias relative to the total domain of interest (Pescott et al., 2019b). Whilst these metrics are often presented in studies whose primary aim is to assess datasets for their limitations, we find that they are rarely presented in studies which use such aggregated species occurrence data to investigate actual patterns of species’ distributions and how they have changed over time [see Ball-Damerow et al. (2019) for a sobering review of the lack of scrutiny where species occurrence data are used across research fields more generally].

One way to encourage the proper use of species occurrence data is to develop software that can facilitate the various tasks involved, thereby easing the burden on researchers’ time. Indeed, a suite of packages have been developed in the R statistical programming environment (R Core Team, 2019) to facilitate the acquisition, cleaning and proper acknowledgement of species occurrence data (Chamberlain et al., 2021; Owens et al., 2021; Zizka et al., 2019). Recently, Zizka et al. (2021) developed what is, to our knowledge, the first R package dedicated to quantifying sampling biases in species occurrence data. The package, called sampbias, quantifies the relative strengths of various geographical biasing factors, such as roads, cities and airports, in a given dataset. While sampbias provides useful information on a set of possible geographical biases in species occurrence data, it is not designed to screen data for biases in other dimensions (e.g. taxonomic, temporal and environmental), and is limited to a specific set of data-biasing mechanisms and the assumption that data point locations are accurate (rather than, for example, grid-based summaries). It would be useful, therefore, to build on the functionality provided by sampbias and develop additional software that can screen species occurrence data for more general biases in a range of possible dimensions. Note that we do not think that bias screening can ever be a completely automatic or easy task: assessing the great number of things that could go wrong, or be misinterpreted, between the numerous data collection, collation, digitisation, and interpretation tasks embodied by the use of any slice of any aggregated database, for any given inferential purpose, should humble any scientist (e.g. Pescott et al., 2018). Nevertheless, making some basic “risk of bias” assessments more straightforward, and raising their profile, is a step in the right direction for ecology.

Here then, we present occAssess: an R package for assessing potential biases in species occurrence data. The package takes a user-supplied dataset and returns a suite of metrics that have been used in the literature to assess species occurrence data for common issues when broad-scale inferences relating to distributions and their changes may be desired. occAssess is designed primarily to assess the suitability of species occurrence data for estimating temporal trends in species’ distributions. Nevertheless, the package should also be useful for those who would like to screen their data for biases of potential importance when estimating spatial variation in species’ occurrences with no explicit reference to time (e.g. using static species distribution models). The aim is to enable quick and easy screening of data for common limitations, thereby enabling researchers to properly scrutinise their data before using it in further analyses, whatever their inferential goal may be. We start by providing an overview of the package, what data it requires, and what outputs it returns. We then provide a worked example using data on the occurrences of leaf-nosed bats and hoverflies in South America over the period 1950 to 2019, and refer the reader to the supporting information where additional vignettes and tutorials can be found. Finally, we discuss different ways in which the package can be used, highlight its limitations, and suggest how it could be improved in the future.

## Package

### Package specifications

occAssess is an open-source R (version >= 4.0.0) package (R Core Team, 2019), built around the existing packages ggplot2 (Wickham, 2016), spatstat (Baddeley et al., 2015), raster (Hijmans, 2019), and stats (R Core Team, 2019). A stable version (1.3.0) can be found at https://github.com/robboyd/occAssess/releases and the development version can be found at https://github.com/robboyd/occAssess.. We provide three vignettes with the package: 1) a tutorial using the data presented in this article; 2) a second example using data that are simulated to be unbiased for the purpose of estimating trends in species’ distributions; and 3) a fully-reproducible example for which all required data are available within the package. Note that all required data is not provided with vignettes one and two; they are provided for instruction, rather than reproducible examples.

### Package structure

occAssess comprises seven discrete functions (Table 1), each of which is designed to assess a common form of potential bias in species occurrence data. The functions each assess species occurrence data in at least one of the spatial, temporal, taxonomic, and environmental dimensions. The user can provide a set of time periods into which the data will be split, meaning that all functions are to some extent temporally explicit. For example, one function assesses spatial bias in the data, but, if multiple periods are specified, then the function provides information on *temporal variation* in spatial bias. We provide the option to split the data into periods to facilitate assessments of the suitability of data for estimating *changes* in species distributions over time. However, in some cases it may be preferable to specify one time period, perhaps covering the entire temporal extent of the data. This may be useful for static species distribution modelling where one simply requires information on, e.g., spatial or environmental bias in the dataset as a whole. At present the time periods must be specified in units of years, and the minimum permitted length for a time period is one year (see Discussion below).

**Table 1.**
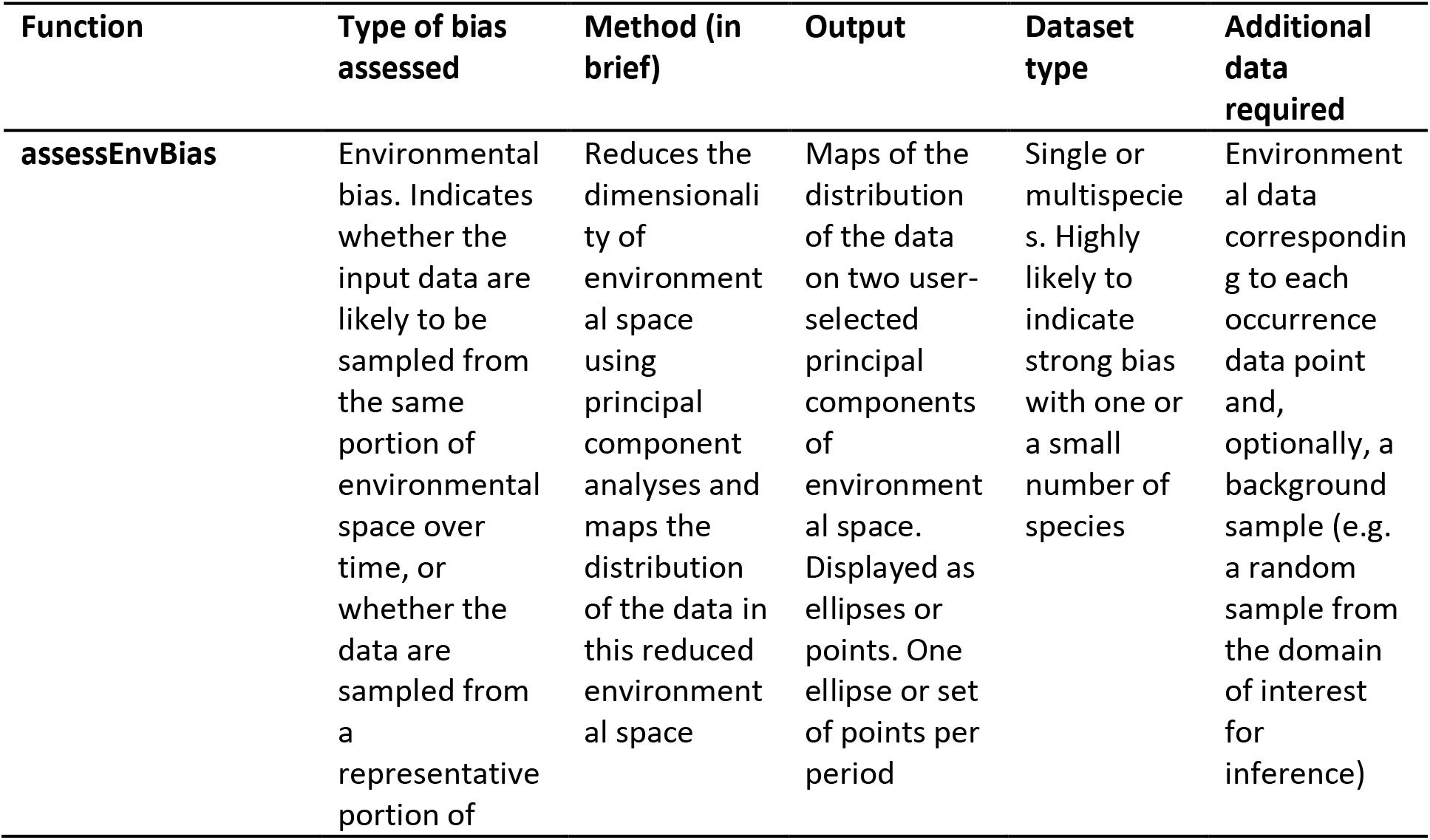
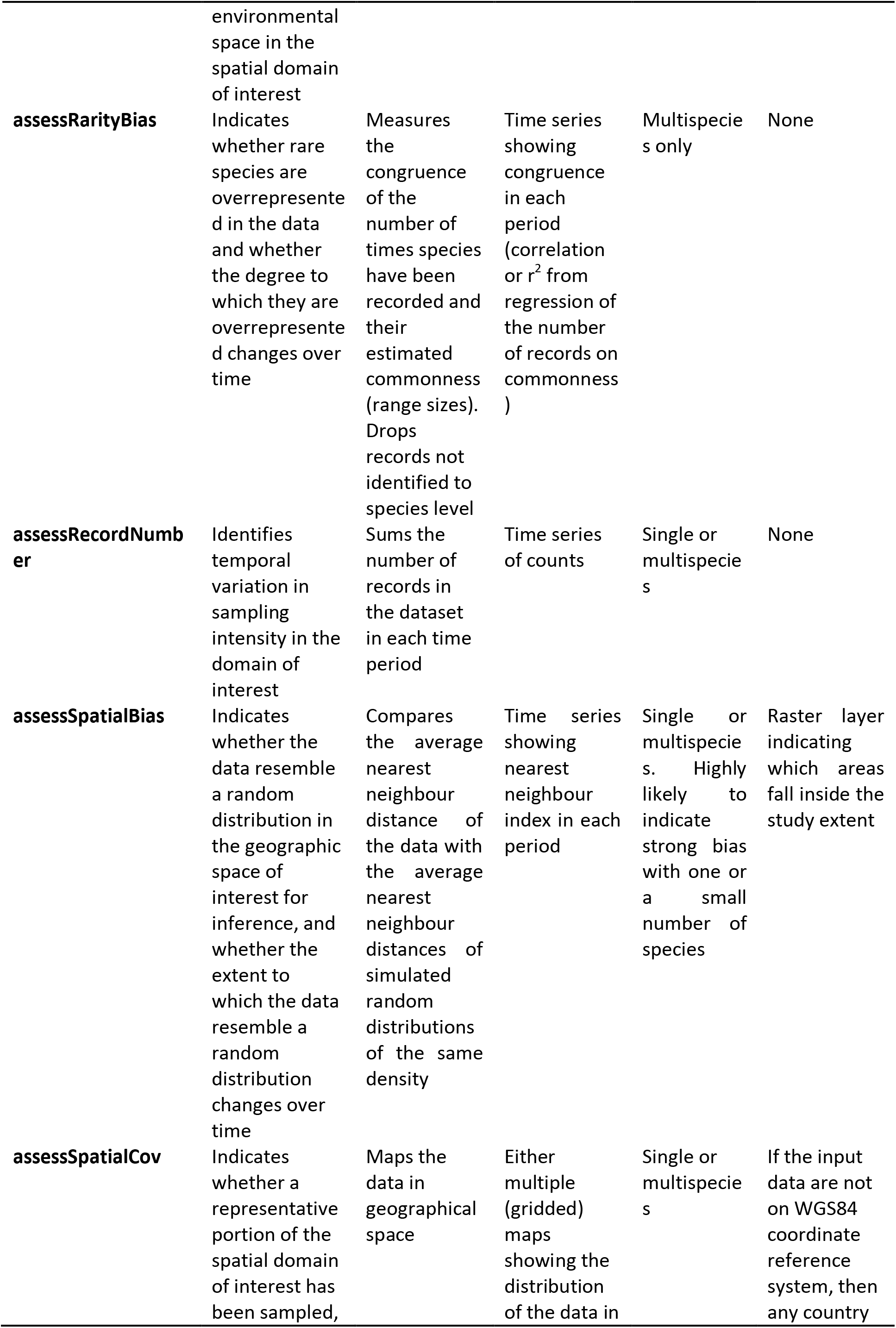
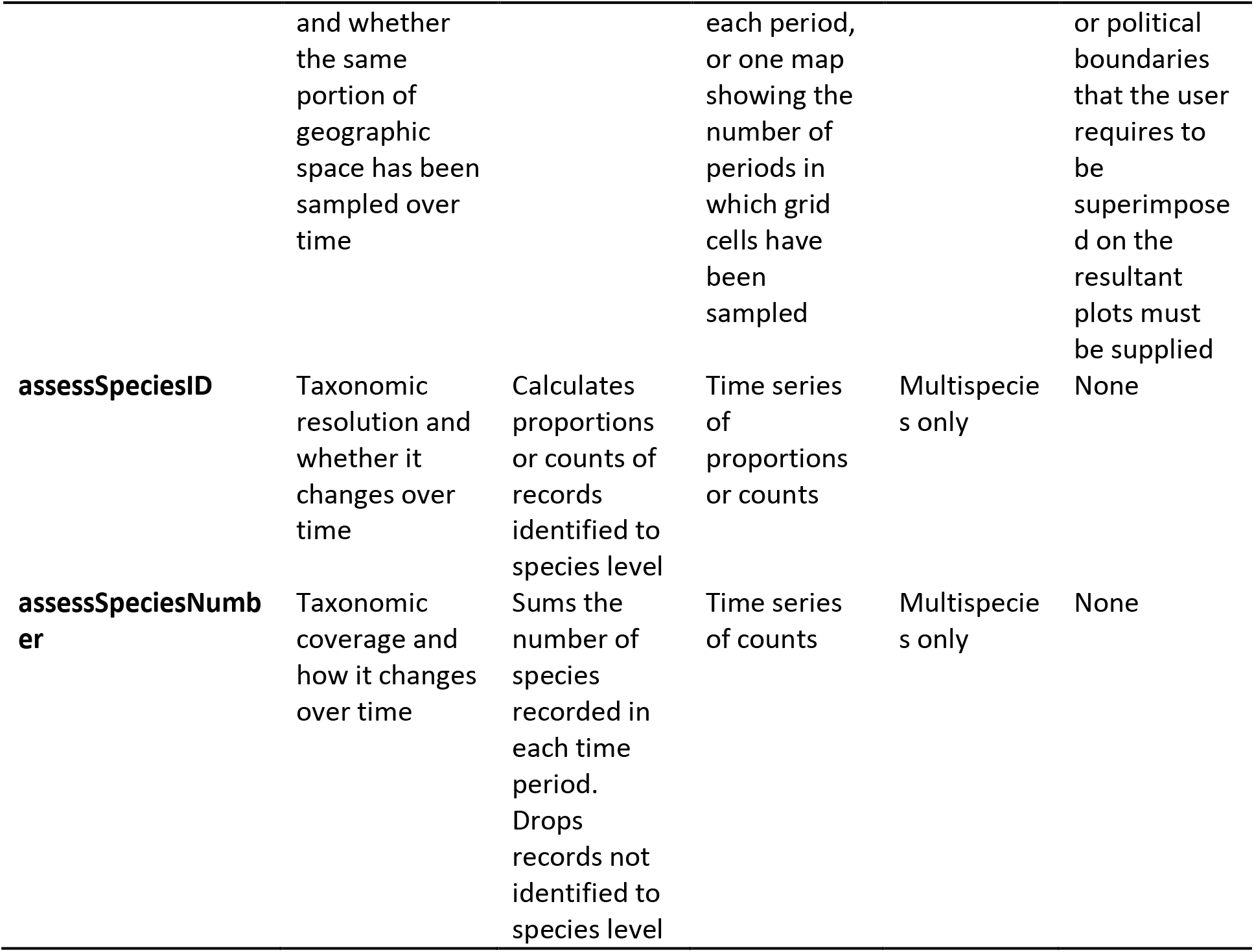
Summary of the functions provided in occAssess. Note that users can opt to split the data into multiple time periods; in this case all functions are temporally explicit and hence provide information on temporal variation in some characteristic of the data. See the worked example in the main text for more details of each function.

#### Input data

For all functions, users must provide their occurrence data and a list of time periods into which the data should be split. The occurrence data must be provided as a dataframe object with six fields: species (species name; note that whilst we use the word “species” here for convenience, essentially any set of taxonomic levels could be used), x (x coordinate), y (y coordinate), spatialUncertainty (uncertainty associated with the x and y coordinates; any units are permitted), year and identifier. The column names in the input data need not match the names of the fields above; rather, the user must pass arguments to each function indicating what columns in their data correspond to which field. This ensures compatibility with data standards such as Darwin Core (https://dwc.tdwg.org/). For example, in Darwin Core, the spatialUncertainty field would be called coordinateUncertaintyInMetres, and the user can provide a mapping by specifying spatialUncertainty = “coordinateUncertaintyInMetres”. We would expect that information on all six required fields would be provided by any typical species occurrence data aggregator, e.g. GBIF. Note that users may specify a threshold spatial uncertainty above which data are dropped before the heuristics are calculated. This allows users to ask the question “how do the biases in my data change if I retain only the more precise records?”. Any coordinate reference system (CRS) may be used. In the spatialUncertainty field, any units are permitted (e.g. metres for eastings/northings, or decimal degrees for lon/lat) but they must be consistent. The identifier field is used to group the data; for example, it may denote specific taxonomic groups, countries, datasets etc. Where there is no information available for a field, its values should be set to NA. See Table 2 for an example set of input data.

**Table 2.** The first six rows of an example dataset as required by occAssess. Note that any units are permitted in the spatialUncertainty field (here metres) but they must be consistent. Also note that the column names in the input data need not match those in this example: users can provide a mapping between their data and the fields presented here using arguments to each function.

| species | x | y | year | spatialUncertainty | identifier |
| --- | --- | --- | --- | --- | --- |
| **Anoura caudifer** | -65.4 | -17.0667 | 1993 | 11839 | Phyllostomidae |
| **Carollia perspicillata** | -65.5497 | -17.1072 | 1993 | 1043 | Phyllostomidae |
| **Carollia perspicillata** | -65.4 | -17.0667 | 1993 | 11839 | Phyllostomidae |
| **Sturnira erythromos** | -65.8692 | -17.2119 | 1993 | 1043 | Phyllostomidae |
| **Platyrrhinus dorsalis** | -65.5497 | -17.1072 | 1993 | 1043 | Phyllostomidae |
| **Artibeus lituratus** | -56 | -25.4667 | 1995 | 11010 | Phyllostomidae |

#### Outputs

Each function returns a list with two elements: a ggplot2 (Wickham, 2016) object, and the data that underpin that plot. The ggplot2 objects generally display the various potential bias metrics for each level of the identifier field (Table 2) and for each time period specified. We provide the outputs as ggplot2 objects because these can be subsequently modified by the user for presentation in e.g. published articles or supplementary material. The functions do not provide any formal recommendation as to whether the data are too biased for any given inferential use; instead, we expect that the heuristics will be used in combination with researchers’ expert judgement to decide on whether mitigating action must be taken, and how this might be done (if indeed it is possible at all). In supplementary material 2 we provide the outputs of occAssess as applied to a simulated dataset that has a random distribution in space and time, and is resolved to species level in all cases; this is taken as an example of a dataset that is unbiased relative to the inferential use case of assessing all species’ distributions in a region over time. These outputs can be used as a point of comparison in that they are likely to provide examples of how the heuristics would appear if a dataset is unbiased.

### Worked example

In this section, we provide a worked example of the functionality of occAssess. We use the package to assess data on the occurrences of leaf-nosed bats and hoverflies in South America over the period 1950 to 2019. The data were downloaded from GBIF (GBIF, 2021; DOI in reference list) and were cleaned for spatial issues (e.g. coordinates matching country centroids, capital cities, biodiversity institutes, etc.) using the CoordinateCleaner package (Zizka et al., 2019). We specify seven time periods, each one decade in duration. We use the identifier field to distinguish between the leaf-nosed bats (Phyllostomidae) and hoverflies (Syrphidae). We do not provide the code in the main text; instead, we refer the reader to the vignette in supplementary material 1 which provides the code for this example. As we introduce each function, and where where applicable, we 1) outline what form of bias it relates to and in what dimension(s); 2) provide the theory behind the metric; 3) indicate where additional inputs – beyond the fields in Table 2 – are required; 4) present the ggplot2 object returned for this case study (noting that the data underpinning these plots are also returned by each function); and 5) give guidance on how to interpret the outputs. We reiterate here that these heuristics are designed to be used alongside expert judgement and careful thought relative to the inferences desired by the analyst – we do not intend any function to provide a simple binary answer to the question “are these data biased for answering my question?”. Biases are challenging!

#### assessRecordNumber()

The simplest function in occAssess, assessRecordNumber, provides a measure of sampling intensity in the domain of interest and how it changes over time (Fig. 1A). Although simple, it is important to understand the extent to which the quantity of data varies over time, because a change in the number of records could reflect a change in recording intensity, which is itself likely to affect the prevalence of particular species in the dataset through time in a non-random fashion (Pescott et al., 2019a).

**Figure 1.**
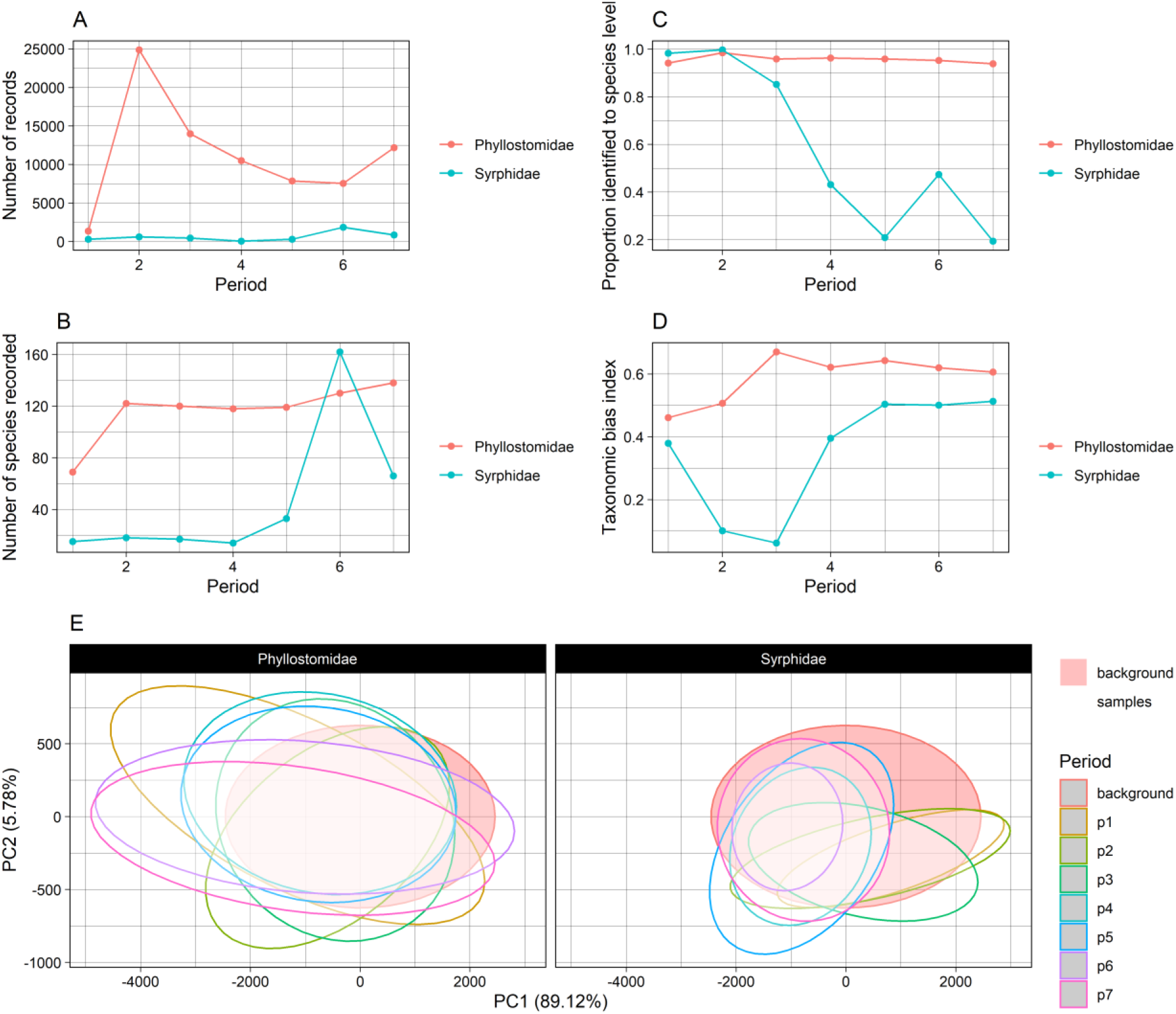
ggplot2 objects returned by A) assessRecordNumber; B) assessSpeciesNumber; C) assessSpeciesID; D) assessRarityBias; and E) assessEnvBias. Note that the ggplot2 objects can be modified by the user (e.g. colours, axis labels, etc.).

One problem that may arise when using assessRecordNumber is that the counts may differ widely between levels of identifier. This can make it difficult to assess temporal variation in record counts for the level(s) with fewer records. To circumvent this problem, we include the option to normalize the counts for each level of identifier. In this case, the indices for each level of identifier fall on comparable scales.

#### assessSpeciesNumber()

The function assessSpeciesNumber returns a measure of taxonomic coverage and how it changes over time. The function sums the number of species recorded in each time period and for each level of identifier and displays the results as time series (Fig. 1B). Of course, changes in the numbers of species recorded could also reflect true extinction/colonisation events in a dataset, but, for heterogeneous, aggregated, data, issues of uneven representativeness across time are considerably more likely. As with assessRecordNumber, users can choose to normalize the species counts for each level of identifier for ease of interpretation.

#### assessSpeciesID()

The function assessSpeciesID provides a measure of taxonomic uncertainty and how it changes over time. By default the function displays the proportion of records identified to species level each year [Fig. 1C, as in Troudet et al. (2018) and Zattara and Aizen (2021)]. Records are considered not identified to species level if they take the value NA. The user has the option to substitute proportions for counts which may be preferable in some circumstances. For example, it is feasible that, due to the increase in the number of records submitted by volunteer citizen scientists over time, the *proportion* of records identified to species level may decrease, but the overall *quantity* may show a different trend.

#### assessRarityBias()

The function assessRarityBias can be used to assess the degree to which rare species are oversampled relative to commoner species, and whether this changes over time. The idea is that, was there no sampling bias, species would be recorded in proportion to their commonness. Commonness can be defined as local abundance or regional occupancy (Gaston, 2011). Following Speed et al. (2018), we define a species’ commonness as the number of grid cells on which it has been recorded – a proxy for regional occupancy. The user may decide on the spatial resolution of the grid cells, and whether commonness is calculated over the entire temporal extent of the data, or separately for each time period (which could have important implications for the interpretation of discovered patterns, given other biases in the dataset).

Once the numbers of times species’ have been recorded, and their commonness, have been calculated, assessRarityBias measures the congruence between these two quantities. The user may decide on one of two methods that the function will use to do this. The first option is to regress the number of records on commonness and use the r^2^ (coefficient of variation) from the fitted model as an indicator of to what extent the number of records are explained by range size. This method is an extension of that used by Barends et al. (2020) and Speed et al. (2018) who fitted analogous regression models and treated each species’ residual as an index of whether they are over- or under-sampled relative to some wider assemblage. This measure ranges from 0, indicating high bias, to 1, indicating low bias. The second option is to use the Pearson’s correlation coefficient between the number of times species’ have been recorded and their commonness as the measure congruence. This measure ranges from −1 – 1, with values closer to 1 indicating smaller bias. Whichever method is chosen, occAssess displays the index for each time period and level of identifier (Fig. 1D). Note that both metrics produced assessRarityBias indicate the strength of the *linear* relationship between range size and the number of records; users may wish to inspect the data for curvillinearity.

#### assessSpatialCov()

The function assessSpatialCov can be used to assess the extent to which the data are spatio-temporally biased; that is, the extent to which the same portion of the geographic domain has been sampled over time — note that this is likely to be crucial for robust estimates of temporal distributional change. The function provides this information in one of two ways (selected by the user). Both methods begin by gridding the data at a user-specified spatial resolution. The first method then returns n ggplot2 objects, where n is the number of levels in the identifier field. Each ggplot2 object contains N maps showing the density of records in each grid cell, where N is the number of time periods. The second method returns one map showing the number of time periods in which each grid cell has been sampled (Figs 2B and 2C; see supplementay materials 2 and 3 for examples using the first method).It is worth pointing out that data originally provided on a grid are often converted to point format by online data aggregators (e.g., using cell centroids). For these data, it is possible that the mismatch between the original grids and the user-specified grid produced by assessSpatialCov could result in some unexpected biases.

**Figure 2.**
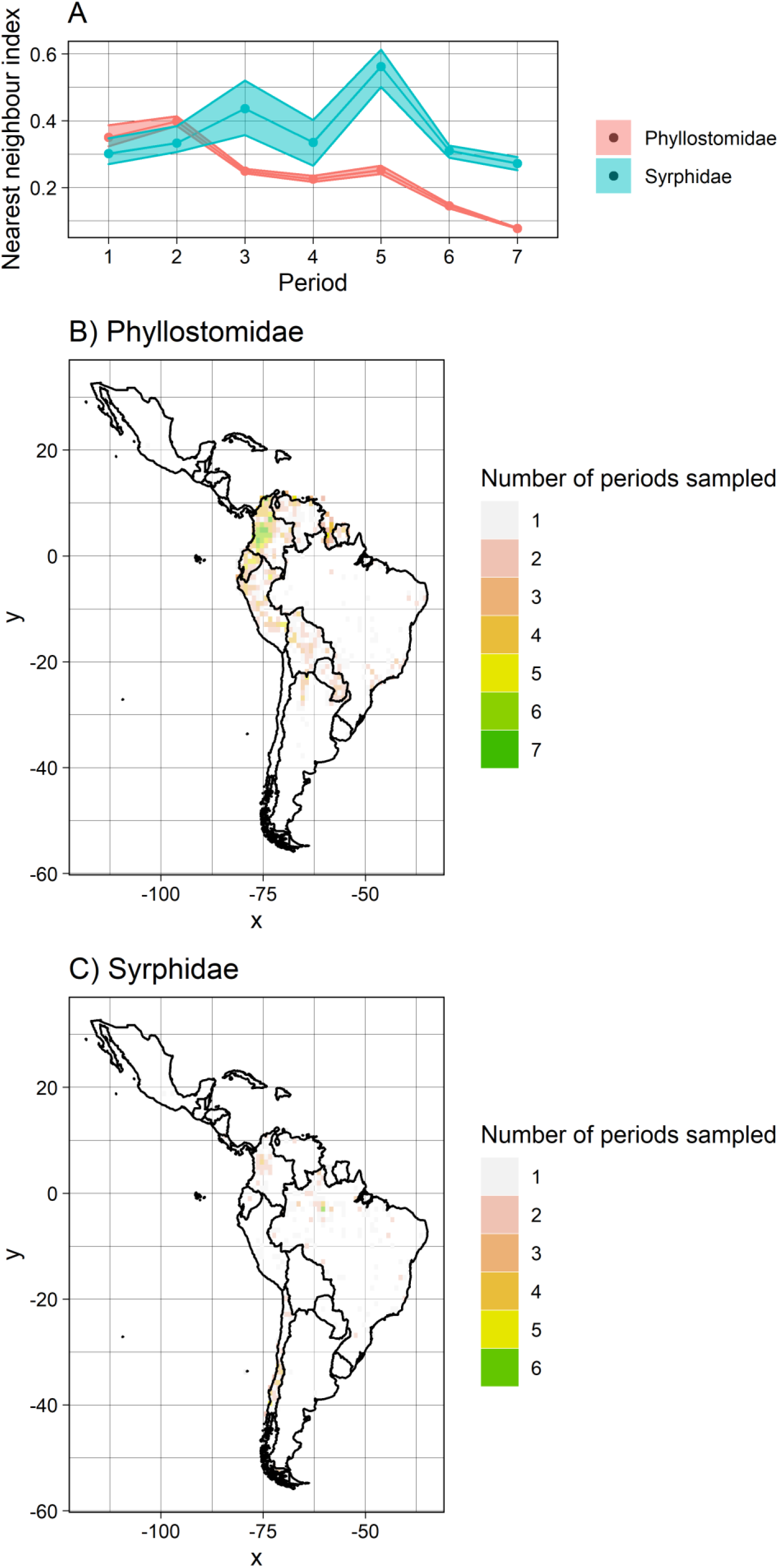
ggplot2 objects returned by assessSpatialBias (A) and assessSpatialCov (B and C). Note that the user can modify these plots (e.g., by changing colours, axis labels, etc.).

In some circumstances users will need to pass additional data to assessSpatialCov to superimpose political/ geographical boundaries on the resultant plots. This is not required where the data are on the WGS84 coordinate reference system; in this case, the user must simply specify the relevant countries, otherwise, a shapefile is required.

#### assessSpatialBias()

The function assessSpatialBias screens data for geographical bias, i.e. the degree to which a sample deviates from a random distribution within the spatial domain of interest. The function is based on the widely-used nearest neighbour index (NNI; Clark and Evans, 1954). The NNI is given as the ratio of the average observed nearest neighbour distances (the Euclidean distance of each data point to its nearest neighbouring point) to the expected average nearest neighbour distance if the data were randomly distributed. In the standard NNI, the average expected nearest neighbour distance for a random distribution is given by 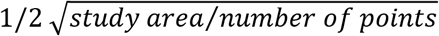. However, in the case of irregularly shaped study boundaries (e.g., political or geographical boundaries), the above formula does not equal the expected average nearest neighbour distances for a random distribution. To circumvent this problem, assessSpatialBias simulates n datasets randomly across the study area in equal number to the occurrence data. The NNI can then be given as the ratio of the average observed nearest neighbour distances to the average of the simulated nearest neighbour distances (Fig. 2A). Another advantage of this approach is that, by simulating n (chosen by the user) random datasets, assessSpatialBias can provide uncertainty associated with the index (the function will display 90% confidence intervals by default). The NNI produced by assessSpatialBias can be interpreted as how far the observed distribution deviates from a random distribution of the same density. Values between 0 and 1 are more clustered than a random distribution, and values between 1 and 2.15 are more widely dispersed (i.e., over-dispersed). See Sumner et al. (2019) for a somewhat similar approach.

It is worth pointing out that the NNI produced by assessSpatialBias is a function of both sampling biases in the data and the true distributions of the focal taxa. If the function is used to assess data for one or a small number of species, the NNI will likely indicate a strong departure from a random distribution. This is to be expected because the geographical distribution of records will reflect e.g., the environmental niche of the target taxa. The function is therefore most appropriate for use with data spanning many species, in which case a more accurate picture of the distribution of sampling is likely to be obtained.

#### assessEnvBias()

The function assessEnvBias can be used to assess species occurrence data for two types of environmental bias: unrepresentative sampling in the environmental space of the domain of interest, and uneven sampling of environmental space over time. The function maps the data in environmental space in each user-specified time period. To do so, additional environmental data are required. As a minimum users must supply environmental data (this can be many variables) at the coordinates of the occurrence data. Users may optionally supply a “background” sample of the same environmental variables; this may be, for example, the environment at random locations across the domain of interest. Whether or not background data are supplied impacts interpretation of the assessEnvBias outputs. If background data are supplied, then the function maps the distribution of the occurrence data in the environmental space of the domain of interest. Otherwise, the data are mapped in the *sampled* environmental space across all periods. In this example we use the standard suite of 19 bioclimatic variables from worldclim (Fick and Hijmans, 2017). These data can be downloaded at https://www.worldclim.org/data/worldclim21.html or through R using the getData function in the raster package (Hijmans et al., 2016).

assessEnvBias reduces the dimensionality of the environmental data using PCAs. It then maps the data in two dimensional environmental space (Fig. 1G), enabling the user to assess whether their data are sampled from the same portion of environmental space across periods or, if background data are supplied, whether the data are sampled from a representative portion of environmental space in the domain of interest. By default, the data are displayed as ellipses delimiting 95% of the occurrence data. Strictly speaking, PCAs assume multivariate normality in the environmental data, and the ellipses displayed by assessEnvBias assume multivariate normality among the principal component scores. Users may wish to assess their data, and the resultant PC scores (which are returned by the function), for normality. If the data are non-normal, then transformations can be applied. If the PC scores are non-normal, it is simple to substitute the ellipses for the actual data points (see s1 for more details). For similar approaches see Pescott et al. (2019b) and Barends et al. (2020). Note that this assessment assumes that the spatial resolution of the environmental data are relevant to the responses of the target organism(s) at the spatial scale of the analysis desired.

## Discussion

In this paper, we have introduced a new R package, occAssess, which enables rapid screening of species occurrence data for biases of potential importance for drawing inferences about species’ distributions and how they have changed over time. The package takes a species occurrence dataset as input, and returns a number of metrics relating to common forms of bias in one or more of the taxonomic, temporal, spatial, and environmental dimensions. None of the metrics provided in the package are new (although some are extended and/or modified). However, we hope that in assembling these metrics in an easy-to-use R package, we will ease the burden on researchers who would like to scrutinise their data. In turn, we hope to promote the proper assessment of species occurrence data *before* they are used in attempts to answer important research questions regarding ecological change. The heuristics returned by occAssess could be provided as, for example, supplementary material to published articles to provide evidence of the fact that a proper assessment has been conducted. In general, we would expect such evidence of assessment to be accompanied by written commentary interpreting the patterns seen and considering their implications for any analyses presented.

We have presented a single example of how occAssess may be used, but it is easy to imagine additional use cases. In our example, we used the identifier field (Table 2) to split the data by taxonomic group (Phyllostomidae and Syrphidae). One might instead use the identifier field to denote specific datasets. For example, one level of identifier could denote a dataset before some newly-digitized data were added, and a second could denote the same data with the addition of the newly-digitized records. It would then be possible to make an assessment of to what extent the data have improved as a result of digitization efforts. occAssess could also be used for model-based data integration (Isaac et al., 2020), where the aim is to exploit the strengths of multiple datasets, each of which could be specified in the identifier field. Another possibility is that occAssess could be used to screen data for single species as opposed to whole taxonomic groups as presented in our worked example. In this case note that some heuristics would require different interpretations; for example, one would expect the data to be biased in environmental space relative to the domain of interest because it would reflect a species’ environmental niche. In summary, we feel that occAssess has the potential to be useful for many applications where species occurrence data are used.

A key feature of occAssess is the periods argument in each function, which enables assessment of how the limitations of a dataset may change over time. We include this feature because a common application of species occurrence data is the estimation of temporal trends in species’ distributions (e.g. Outhwaite et al., 2019; Pescott et al., 2019a; Powney et al., 2019). For some applications, however, it may be more appropriate to consider an entire dataset as comprising one time period, thereby removing the temporal dimension. An obvious example is where data are to be used for species distribution modelling (SDM). In this case the objective is typically estimation of spatial variation in species’ occurrences with no explicit reference to time (Guisan, 2017). Where occAssess is used to screen data for use in SDMs, we suggest that the functions relating to spatial and environmental bias will be of most importance, namely assessSpatialBias, assessSpatialCov and assessEnvBias (although the other functions could still provide important context on the temporal dynamics of the dataset).

The functions in occAssess provide heuristics relating to the quality of species occurrence data, but stop short of making a formal recommendation as to whether the data are of sufficient quality for any given use. It would not be appropriate to provide such recommendations, because the utility of species occurrence data depend not only their biases, but also on the question being asked and the methods used to answer it. For example, it may be possible to obtain relatively unbiased predictions of species’ geographical distributions using SDMs, even when the data themselves are spatially and environmentally biased. Phillips et al. (2009) developed the “target group” approach whereby background data are generated with similar sampling biases to the occurrence data. This approach helps SDMs to distinguish between suitable and unsuitable habitats as opposed to popular and unpopular sampling locations. There have also been attempts to correct for changes in recorder effort statistically, thereby enabling estimation of how species’ distributions have changed over time from biased data (Franklin, 1999; Hill, 2012; Isaac et al., 2014; Szabo et al., 2010; Telfer et al., 2002; Van Strien et al., 2013). While it is not always clear to what extent the above-mentioned methods achieve the goal of mitigating for sampling biases, the point remains that relatively informative inferences may be possible from biased data where the biases can either be modelled, or reduced through appropriate resolution-based aggregation (Pescott et al., 2019a), or through more complex methods designed to leverage unbiased estimates of model parameters from additional probability samples (e.g. Ahmad Suhaimi et al., 2021). It is for this reason that we suggest the metrics provided by occAssess be consulted in combination with other relevant information in order to decide whether or not a dataset is of sufficient quality for use for a given inferential purpose.

The version of occAssess presented here has two key limitations. First, the temporal unit is the year, meaning that the package can say nothing about intra-annual biases (e.g. phenological patterns in the data). In future versions it might be feasible to increase the temporal resolution of the package. Second, it will not always be possible to tease apart biases from true biological phenomena using the package alone. For example, assessSpatialBias indicates whether the data deviate from a random distribution but, particularly where there are few species in the dataset, it might not be clear whether this reflects sampling biases or species’ true distributions. To disentangle sampling biases and the biological truth it will always be preferable to solicit advice from experts who are familiar with the biology of the focal taxa – we stress again that our package is a compliment to, not substitute for, expert knowledge.

## Supporting information

vignette 1

vignette 2

vignette 3

## Acknowledgements

We would like to thank Francesca Mancini, Nick Isaac and Robert Cooke for their helpful comments on the heuristics presented in this article. RB, GP and CC were funded by the SURPASS2 project under the Newton Fund Latin America Biodiversity Programme: Biodiversity - Ecosystem services for sustainable development, awarded by the UKRI Natural Environment Research Council (NERC) NE/S011870/2. We thank SURPASS2 partners Francisco Fonturbel, Marcelo Aizen, Eduardo Zattara, Antonio Saraiva and Jeff Ollerton for comments on the outputs presented. The contribution of OLP was supported by the Natural Environment Research Council award number NE/R016429/1 as part of the UK Status, Change and Projections of the Environment (UK-SCAPE) programme delivering National Capability.

## Supporting information

We provide three vignettes as supporting information: S1) a vignette showing the code for the worked example presented in the manuscript; S2) a second example using data that have been simulated to unbiased relative to the inferential goal of estimating temporal trends in species’ distributions; and 3) a fully-reproducible example for which all data are available in the package.

