## Supplementary material for "*occAssess*: An R package for assessing potential biases in species occurrence data": vignette 1

occAssess

if (!"occAssess" %in% installed.packages()) devtools::install_github("https://github.com/robboyd/occAssess")
library(occAssess)

### Introduction

### What does occAssess do?

occAssess enables quick and easy screening of species occurrence data for common forms of bias.

#### How does it work?

The package comprises a number of discrete functions, each of which is designed to assess a common form of bias. Generally speaking, users must pass three bits of information to the functions: 1) their occurrence data to the functions; 2) and list of time periods into which the resulting metrics will be split; and 3) the names of the columns in their data that correspond to the required fields in occAssess. Ouputs are generally provided in list format, with one element containing a ggplot2 object, and a second containing the data that underpins that plot.

#### A worked example

In this document I demonstrate the utility of occAssess by applying it to data on flower-visiting taxa occurrences in South America over the period 1950 to 2019. The data were extracted from GBIF and can be accessed at <https://doi.org/10.15468/dl.wqz6z3>.

##### Inputs

###### Occurrence data

occAssess requires a data.frame with the following fields: species (species name), x (x coordinae of record), y (y coordinate), year, spatialUncertainty (how much uncertainty is associated with the x and y coordinates; units do not matter but must be consistent) and identifier. The identifier is used to split the data into groups; for example, it could represent taxonomic groups or specific datasets. Where you have no information for a field, its value should be set to NA. The column names in your data need not match the fields needed for occAssess; instead, you must specify which columns corresponds to which of the required fields. This enables compatability with data standards such as Darwin core.

```

spDat <- read.csv("C:/Users/Rob.Lenovo-PC/Documents/surpass/Data/GBIF/07.04.21/cleanedData.csv")

str(spDat)
#> 'data.frame':    82710 obs. of  6 variables:
#>  $ species           : chr  "Artibeus lituratus" "Carollia perspicillata" "Sturnira parvidens" "Sturnira tildae" ...
#>  $ x                 : num  -75.1 -75.1 -67.8 -61.4 -72 ...
#>  $ y                 : num  5.27 5.28 10.4 5.98 7.32 ...
#>  $ year              : int  2019 2019 1960 1966 1968 1968 1968 1986 1981 1967 ...
#>  $ spatialUncertainty: num  NA NA NA NA NA NA NA NA NA NA ...
#>  $ identifier        : chr  "Phyllostomidae" "Phyllostomidae" "Phyllostomidae" "Phyllostomidae" ...
```

###### Periods

In addition to your occurrence data, you must also provide a number of periods into which your data will be split (thi can be one period covering e.g. your whole study extent). In this example I will split the data into decades over the period 1950 to 2019. Periods are defined as follows:

periods <- list(1950:1959, 1960:1969, 1970:1979, 1980:1989, 1990:1999, 2000:2009, 2010:2019)

##### Functions

All of the functions in occAssess require two common arguments (in addition to the arguments that specify which column matches which field): dat and periods (outlined above). I will run through each function in the following, indicating where additional arguments are required. Generally, the functions in occAssess return a list with two elements: one being a ggplot2 object, with a separate panel or group for each level of identifier; and a second with the data underpinning the plot.

###### assessRecordNumber

The first function I will introduce is the simplest: assessRecordNumber. This function simply plots out the number of records per year in your dataset.

```

nRec <- assessRecordNumber(dat = spDat,
                           periods = periods,
                           species = "species",
                           x = "x",
                           y = "y",
                           year = "year",
                           spatialUncertainty = "spatialUncertainty",
                           identifier = "identifier",
                           normalize = FALSE)

str(nRec$data)
#> 'data.frame':    14 obs. of  3 variables:
#>  $ val   : int  1363 24855 13971 10499 7843 7546 12183 302 613 461 ...
#>  $ group : chr  "Phyllostomidae" "Phyllostomidae" "Phyllostomidae" "Phyllostomidae" ...
#>  $ Period: int  1 2 3 4 5 6 7 1 2 3 ...

nRec$plot
```

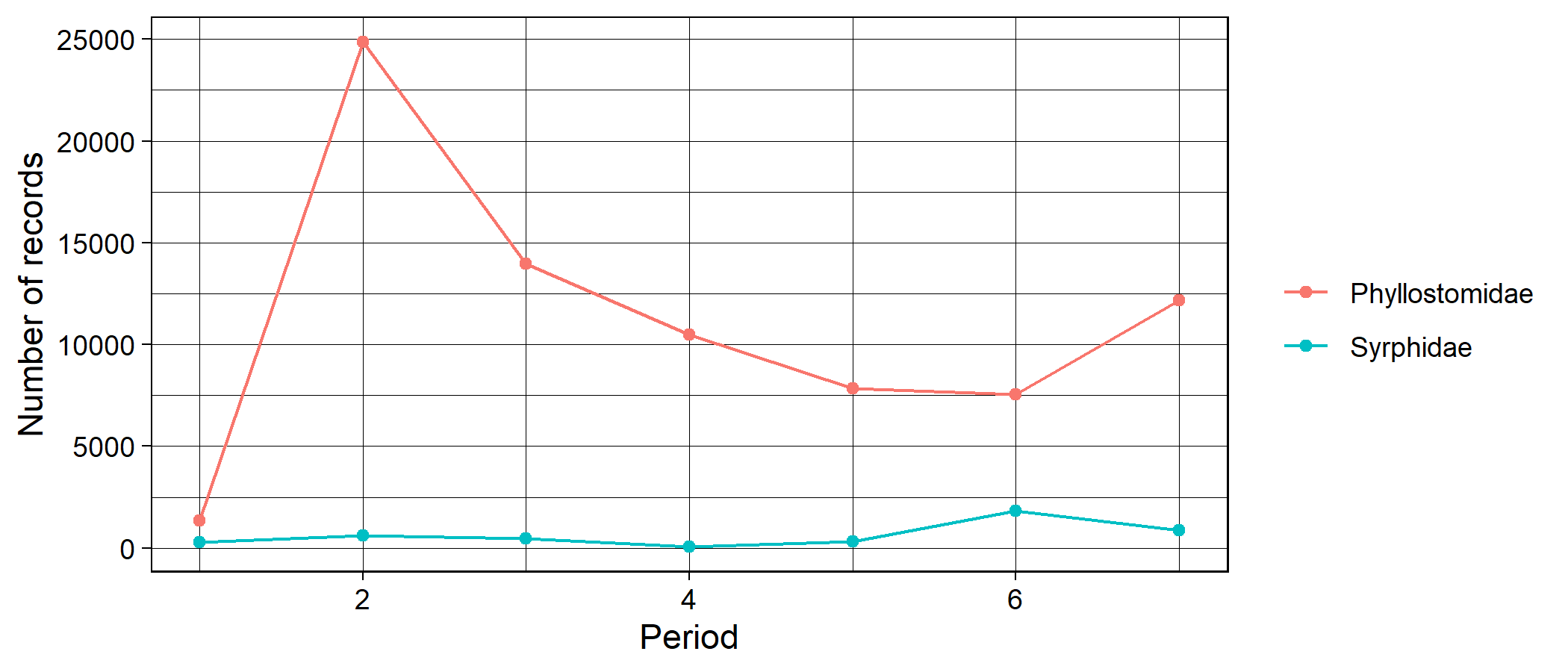

plot of chunk unnamed-chunk-4

This function enables researchers to quickly establish how the number of records has changed over time. Note the argument “normalize” which, if TRUE, will rescale the counts for each level of identifier to enable comparisons where they are very different.

###### assessSpeciesNumber

In addition to the number of records, you may wish to know how the number of species (taxonomic coverage) in your dataset changes over time. For this you can use the function assessSpeciesNumber:

```

nSpec <- assessSpeciesNumber(dat = spDat,
                             periods = periods,
                             species = "species",
                             x = "x",
                             y = "y",
                             year = "year",
                             spatialUncertainty = "spatialUncertainty",
                             identifier = "identifier",
                             normalize = FALSE)

str(nSpec$data)
#> 'data.frame':    14 obs. of  3 variables:
#>  $ val   : int  69 122 120 118 119 130 138 15 18 17 ...
#>  $ group : chr  "Phyllostomidae" "Phyllostomidae" "Phyllostomidae" "Phyllostomidae" ...
#>  $ Period: int  1 2 3 4 5 6 7 1 2 3 ...

nSpec$plot
```

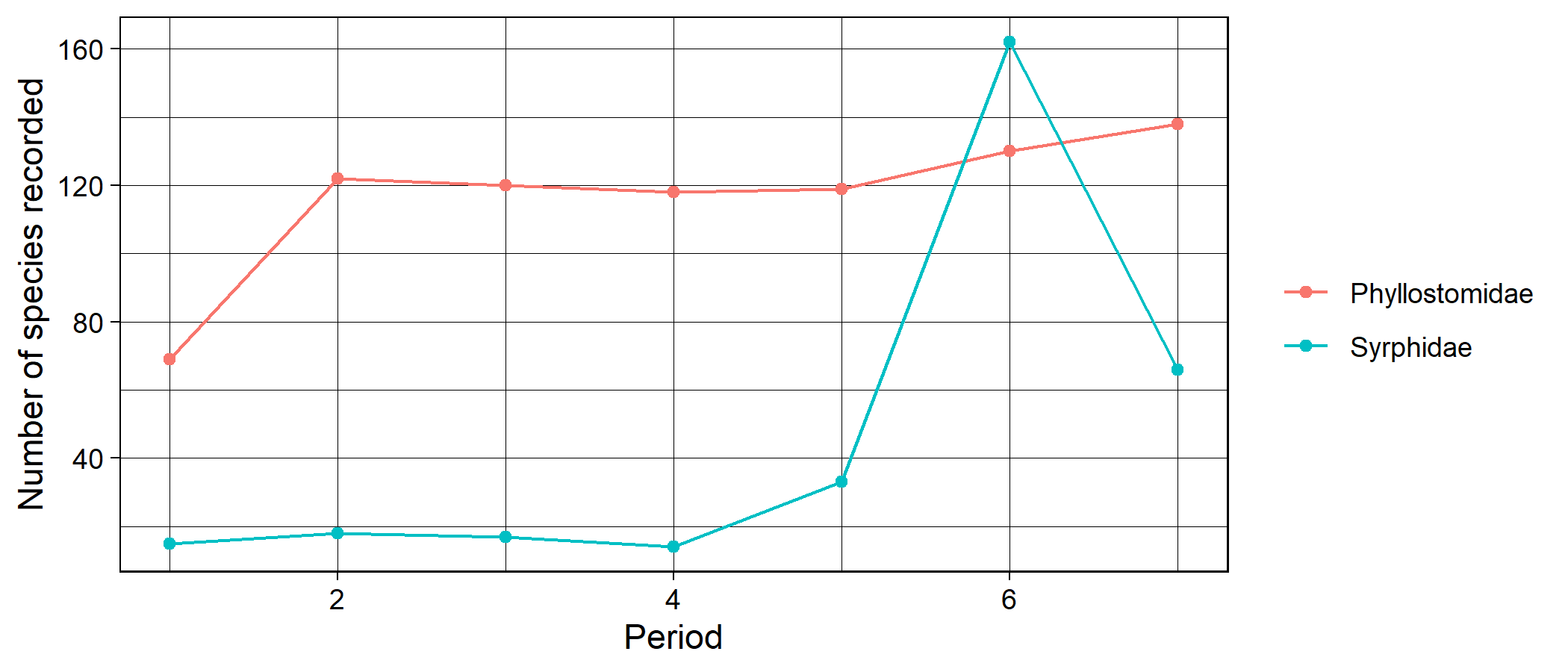

plot of chunk unnamed-chunk-5

###### assessSpeciesID

It has been speculated that apparent changes in taxonomic coverage could, in fact, reflect a change in taxonomic expertise over time. For example, if fewer individuals have the skill to identify certain species, then it may not appear in your dataset in the later periods. The function assessSpeciesID treats the proportion of species identified to species level as a proxy for taxonomic expertise:

```

propID <- assessSpeciesID(dat = spDat,
                           periods = periods,
                           type = "proportion",
                           species = "species",
                           x = "x",
                           y = "y",
                           year = "year",
                           spatialUncertainty = "spatialUncertainty",
                           identifier = "identifier")

str(propID$data)
#> 'data.frame':    14 obs. of  3 variables:
#>  $ prop  : num  0.941 0.986 0.959 0.963 0.959 ...
#>  $ group : chr  "Phyllostomidae" "Phyllostomidae" "Phyllostomidae" "Phyllostomidae" ...
#>  $ Period: int  1 2 3 4 5 6 7 1 2 3 ...

propID$plot
```

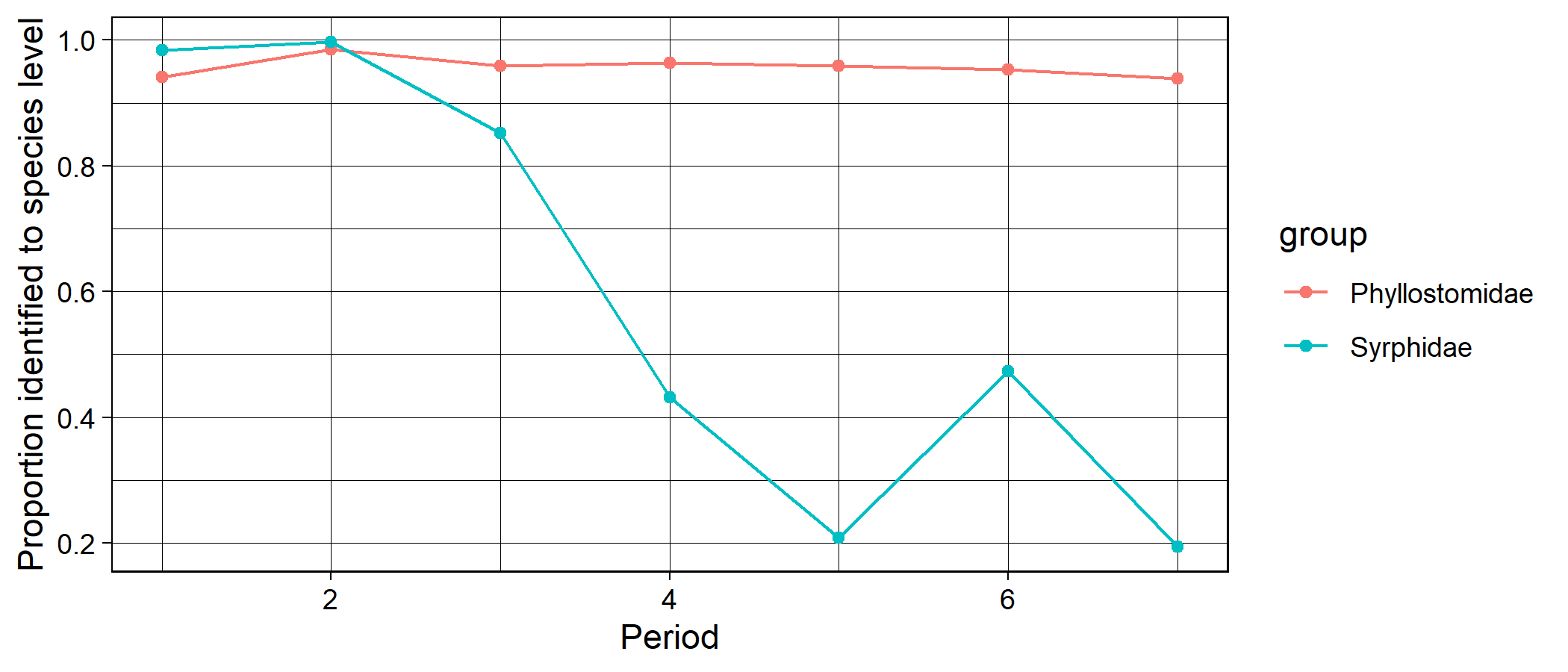

plot of chunk unnamed-chunk-6

The argument “type” can take the values proportion (proportion of records identified to species level) or count (number of records identified to species level).

###### assessRarityBias

A number of studies have defined taxonomic bias in a dataset as the degree of proportionality between species’ range sizes (usually proxied by the number of grid cells on which it has been recorded) and the total number of records. One can regress the number of records on range size, and the residuals give an index of how over-or undersampled a species is given its prevalence. The function assessSpeciesBias conducts these analyses for each time period, and uses the r2 value from the linear regressions as an index proportionality between range sizes and number of records. Higher values indicate that species’ are sampled in proportion to their range sizes whereas lower values indicate that some species are over- or undersampled.

```

taxBias <- assessRarityBias(dat = spDat,
                            periods = periods,
                            res = 0.5,
                            prevPerPeriod = FALSE,
                            species = "species",
                            x = "x",
                            y = "y",
                            year = "year",
                            spatialUncertainty = "spatialUncertainty",
                            identifier = "identifier")
#> Warning in assessRarityBias(dat = spDat, periods = periods, res = 0.5, prevPerPeriod = FALSE, : Removing 4843 records
#> because they are do not identified to species level.

str(taxBias$data)
#> 'data.frame':    14 obs. of  3 variables:
#>  $ period: int  1 2 3 4 5 6 7 1 2 3 ...
#>  $ id    : chr  "Phyllostomidae" "Phyllostomidae" "Phyllostomidae" "Phyllostomidae" ...
#>  $ index : num  0.459 0.562 0.716 0.688 0.707 ...

taxBias$plot +ggplot2::ylim(c(0,1))
```

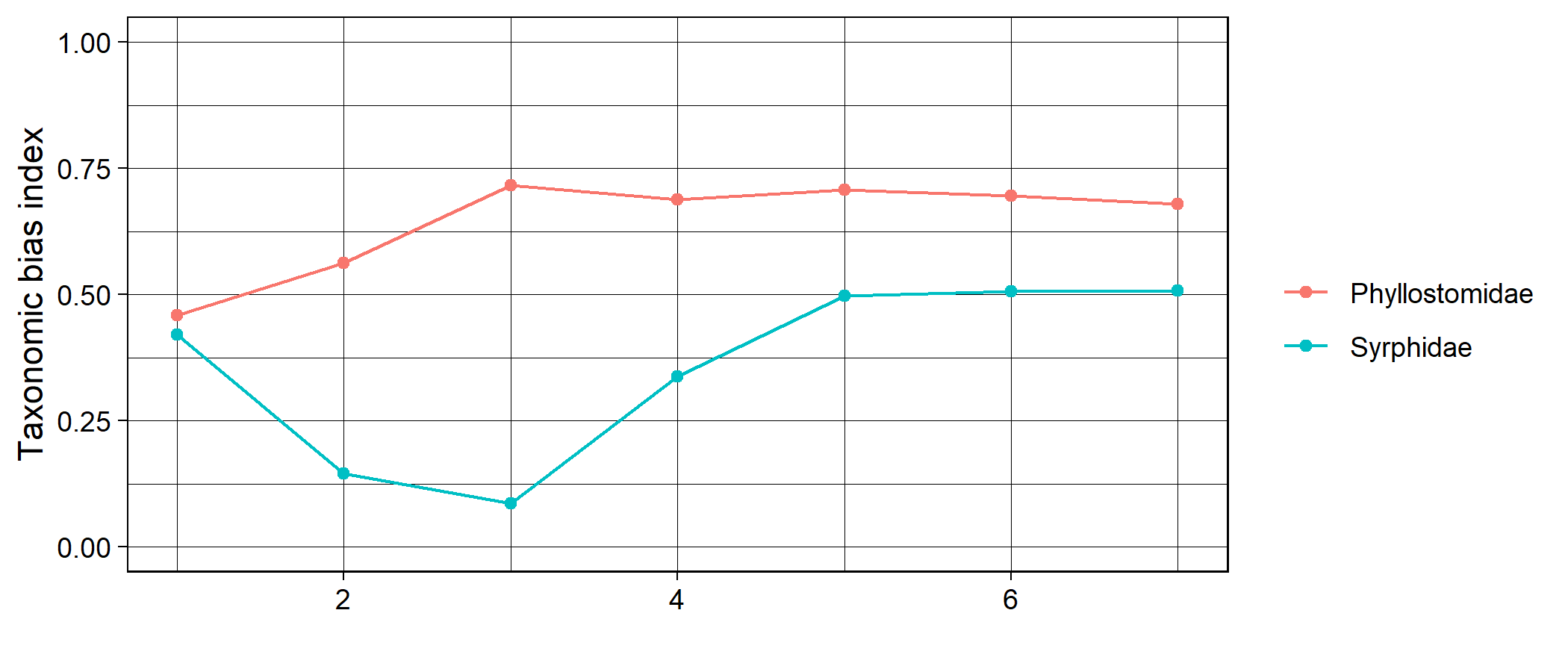

plot of chunk unnamed-chunk-7

###### assessSpatialCov

The function assessSpatialCov grids your data at a specified spatial resolution then maps it in geographic space:

```

maps <- assessSpatialCov(dat = spDat,
                         periods = periods,
                         res = 0.5,
                         logCount = TRUE,
                         countries = c("Brazil", "Argentina", "Chile", "Colombia", "Ecuador", "Bolivia", "Uruguay",
                         "Venezuela",  "Paraguay", "Peru", "Suriname", "Mexico", "Panama", "Costa Rica", "Honduras",
                         "Nicaragua", "Belize", "El Salvador", "Guatemala", "Cuba", "Dominican Republic"),
                         species = "species",
                         x = "x",
                         y = "y",
                         year = "year",
                         spatialUncertainty = "spatialUncertainty",
                         identifier = "identifier")

maps[[1]] + ggplot2::xlim(c(-90, -30)) # Phyllostomidae
#> Warning: Removed 47859 rows containing missing values (geom_tile).
```

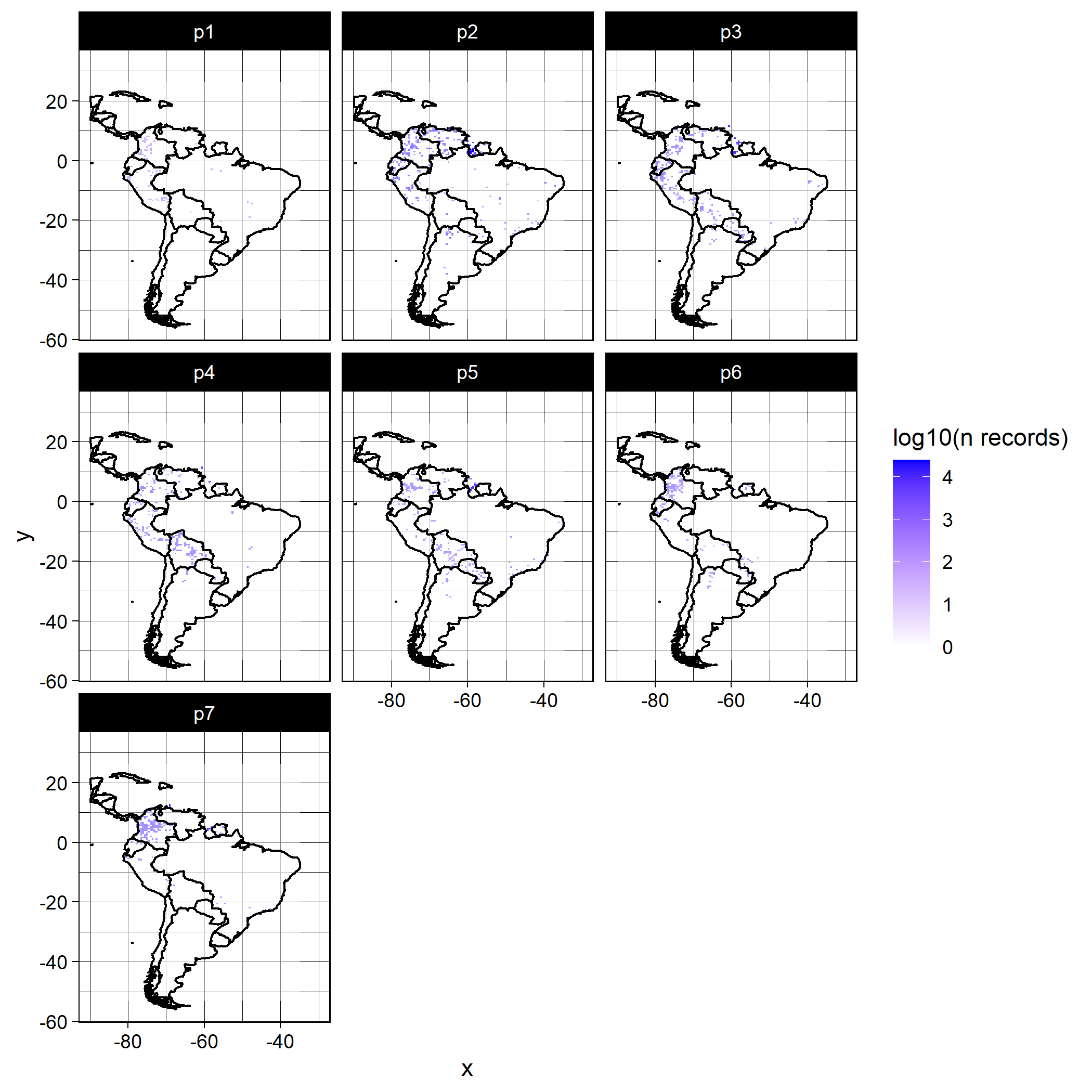

plot of chunk unnamed-chunk-8

```

maps[[2]] + ggplot2::xlim(c(-90, -30)) # Syrphidae
#> Warning: Removed 47859 rows containing missing values (geom_tile).
```

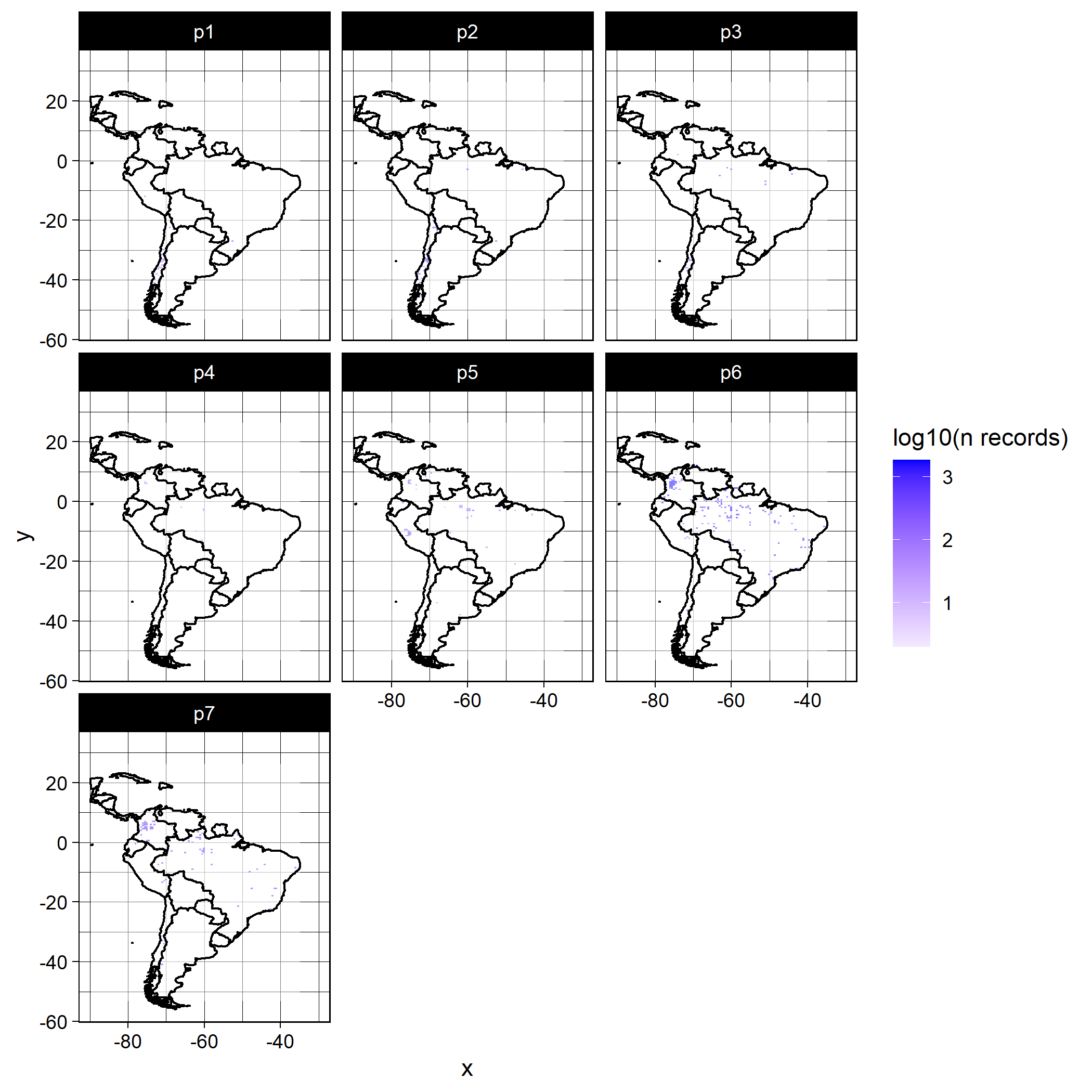

plot of chunk unnamed-chunk-8

As you can see there are three new arguments to be specified. res is the spatial resolution at which you would like to map the data (units depend on you coordinate reference system, e.g. m if easting and northing, and decimal degress in lon/ lat); logCount indicates whether or not you would like to log10 transform the counts for visual purposes; and countries defines the countries covered by your data. Countries must be specified in order to plot their boundaries.

###### assessSpatialBias

Even if your data has good spatial coverage, it may be biased; that is to say, it may deviate from a random distribution in space. The function assessSpatialBias provides an index of how far your data deviates from a random distribution. To do this is simulates an equal number of points to your data randomly across your study region. Then, for each time period, it calculates the average nearest neighbour distance across your data points and divides it by the average nearest neighbour distance from the random sample. If the index is lower than one then your data is more clustered than the random sample, and if it is above one it is more dispersed. To delineate your study area, you must provide a mask layer. The mask is a raster object which is has numeric values within your study area, and is NA outside of your study area. Here, I’ll use some climate data from South America as a mask layer:

```

mask <- raster::raster("C:/Users/Rob.Lenovo-PC/Documents/surpass/Data/occAssessData/climSA.asc")

mask
#> class      : RasterLayer
#> dimensions : 410, 279, 114390  (nrow, ncol, ncell)
#> resolution : 0.1666667, 0.1666667  (x, y)
#> extent     : -81.33333, -34.83333, -55.83333, 12.5  (xmin, xmax, ymin, ymax)
#> crs        : NA
#> source     : climSA.asc
#> names      : climSA

spatBias <- assessSpatialBias(dat = spDat,
                              periods = periods,
                              mask = mask,
                              nSamps = 10,
                              degrade = TRUE,
                              species = "species",
                              x = "x",
                              y = "y",
                              year = "year",
                              spatialUncertainty = "spatialUncertainty",
                              identifier = "identifier")
#> Registered S3 method overwritten by 'spatstat.geom':
#>   method     from
#>   print.boxx cli
#> Warning in FUN(X[[i]], ...): Fewer than 100 records in period 1 for Syrphidae . View this result with caution.
#> Warning in FUN(X[[i]], ...): Fewer than 100 records in period 2 for Syrphidae . View this result with caution.
#> Warning in FUN(X[[i]], ...): Fewer than 100 records in period 3 for Syrphidae . View this result with caution.
#> Warning in FUN(X[[i]], ...): Fewer than 100 records in period 4 for Syrphidae . View this result with caution.
#> Warning in FUN(X[[i]], ...): Fewer than 100 records in period 5 for Syrphidae . View this result with caution.
#> Warning: `guides(<scale> = FALSE)` is deprecated. Please use `guides(<scale> = "none")` instead.

str(spatBias$data)
#> 'data.frame':    14 obs. of  5 variables:
#>  $ mean      : num  0.397 0.437 0.278 0.247 0.277 ...
#>  $ upper     : num  0.424 0.449 0.288 0.257 0.289 ...
#>  $ lower     : num  0.377 0.416 0.272 0.234 0.266 ...
#>  $ Period    : chr  "1" "2" "3" "4" ...
#>  $ identifier: chr  "Phyllostomidae" "Phyllostomidae" "Phyllostomidae" "Phyllostomidae" ...

spatBias$plot + ggplot2::ylim(c(0,1))
```

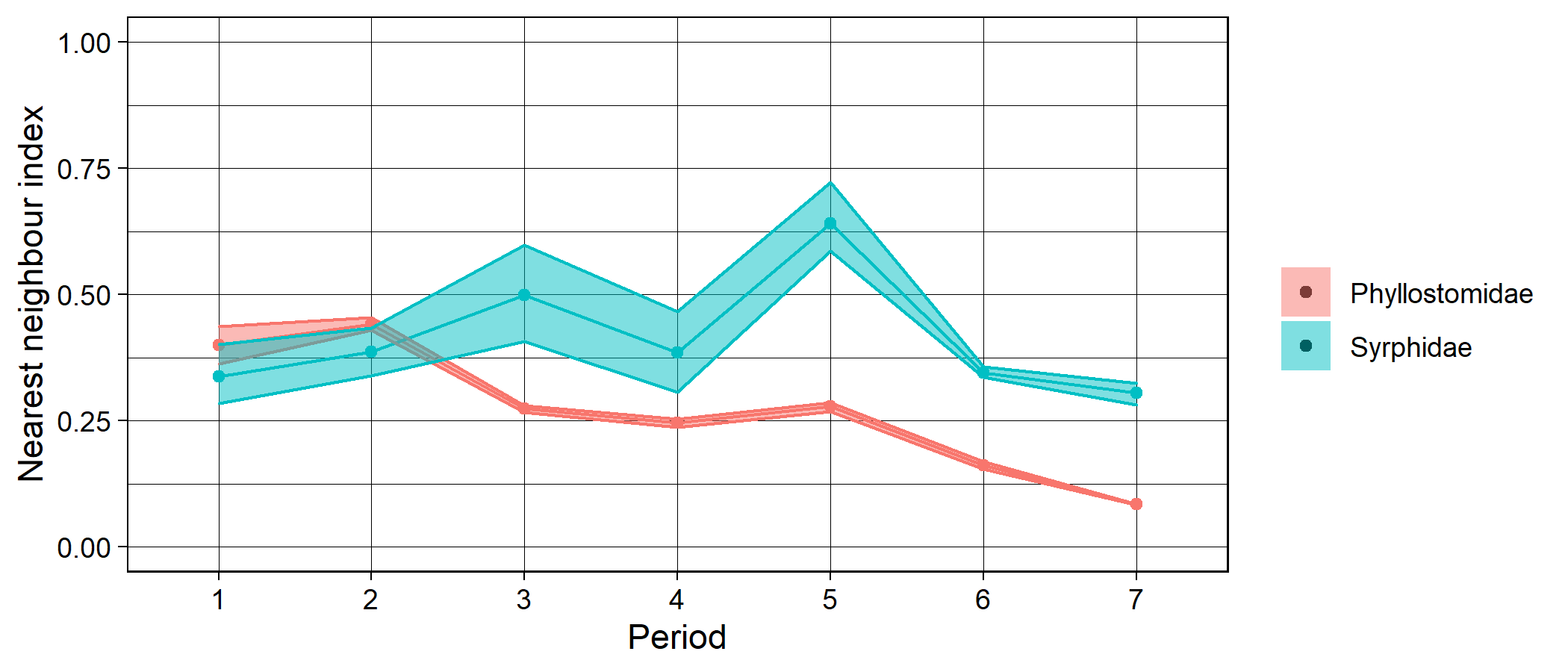

plot of chunk unnamed-chunk-10

The argument nSamps indicates how many random distributions should be drawn, and the argument degrade = TRUE indicates that any duplicated coordinates within a time period and for a given level of identifier are removed. The shaded regions on the plot indicate the 5th and 95th percentiles of the nearest neighbour index calculated over nSamps random samples.

###### assessEnvBias

Spatial bias in your dataset does not necessarily tell you anything about environmental bias. The function assessEnvBias assesses the degree to which your data are biased across time periods in environmental space. To do this we first need to get some climate data. I will use the standard suite of 19 bioclimatic variables from worldclim. It is possible to get this data through R using the raster package, but here I will use my local version for speed. The environmental data for assessEnvBias can be processed as follows:

```
## How to get the data using raster::getData()

#clim <- raster::getData("worldclim",var="bio",res=10)

clim <- raster::stack(list.files("C:/Users/Rob.Lenovo-PC/Documents/surpass/Data/bio/bio/",
                           full.names = T)) # my local version (19 raster layers)

# delineate south america in the climate data

shp <- raster::shapefile("C:/Users/Rob.Lenovo-PC/Documents/surpass/Data/South America country boundaries/South America country boundaries/data/commondata/data0/southamerica_adm0.shp")
#> Warning in OGRSpatialRef(dsn, layer, morphFromESRI = morphFromESRI, dumpSRS = dumpSRS, : Discarded ellps WGS 84 in Proj4
#> definition: +proj=merc +a=6378137 +b=6378137 +lat_ts=0 +lon_0=0 +x_0=0 +y_0=0 +k=1 +units=m +nadgrids=@null +wktext
#> +no_defs
#> Warning in OGRSpatialRef(dsn, layer, morphFromESRI = morphFromESRI, dumpSRS = dumpSRS, : Discarded datum WGS_1984 in
#> Proj4 definition: +proj=merc +a=6378137 +b=6378137 +lat_ts=0 +lon_0=0 +x_0=0 +y_0=0 +k=1 +units=m +nadgrids=@null +wktext
#> +no_defs
#> Warning in showSRID(wkt2, "PROJ"): Discarded ellps WGS 84 in Proj4 definition: +proj=merc +a=6378137 +b=6378137 +lat_ts=0
#> +lon_0=0 +x_0=0 +y_0=0 +k=1 +units=m +nadgrids=@null +wktext +no_defs +type=crs
#> Warning in showSRID(wkt2, "PROJ"): Discarded datum World Geodetic System 1984 in Proj4 definition

shp <- sp::spTransform(shp, raster::crs(clim))

clim <- raster::crop(clim, raster::extent(shp))

clim <- raster::mask(clim, shp)

## exract climate data at coordinates of occurrence data

envDat <- raster::extract(clim, spDat[, c("x", "y")])

## extract background environmental data

backgroundEnvDat <- raster::sampleRandom(clim, size = 10000,
                       xy = F)
```

assessEnvBias conducts a principal component analysis on your environmental data, then maps your occurrence data in 2d environmental space. If backgroundEnvDat is supplied, then the PCA is conducted on this sample, and the occurrence data are projected on to this plane. Otherwise, the PCA is conducted on the environmental data corresponding to the occurrence data and there is no reference to background environmental space.

```

envBias <- assessEnvBias(dat = spDat,
                         species = "species",
                         x = "x",
                         y = "y",
                         year = "year",
                         spatialUncertainty = "spatialUncertainty",
                         identifier = "identifier",
                         envDat = envDat,
                         backgroundEnvDat = backgroundEnvDat,
                         xPC = 1,
                         yPC = 2,
                         periods = periods) # xPC and yPC indicate which principal components to set as the x and y axes,respectively


envBias$plot
```

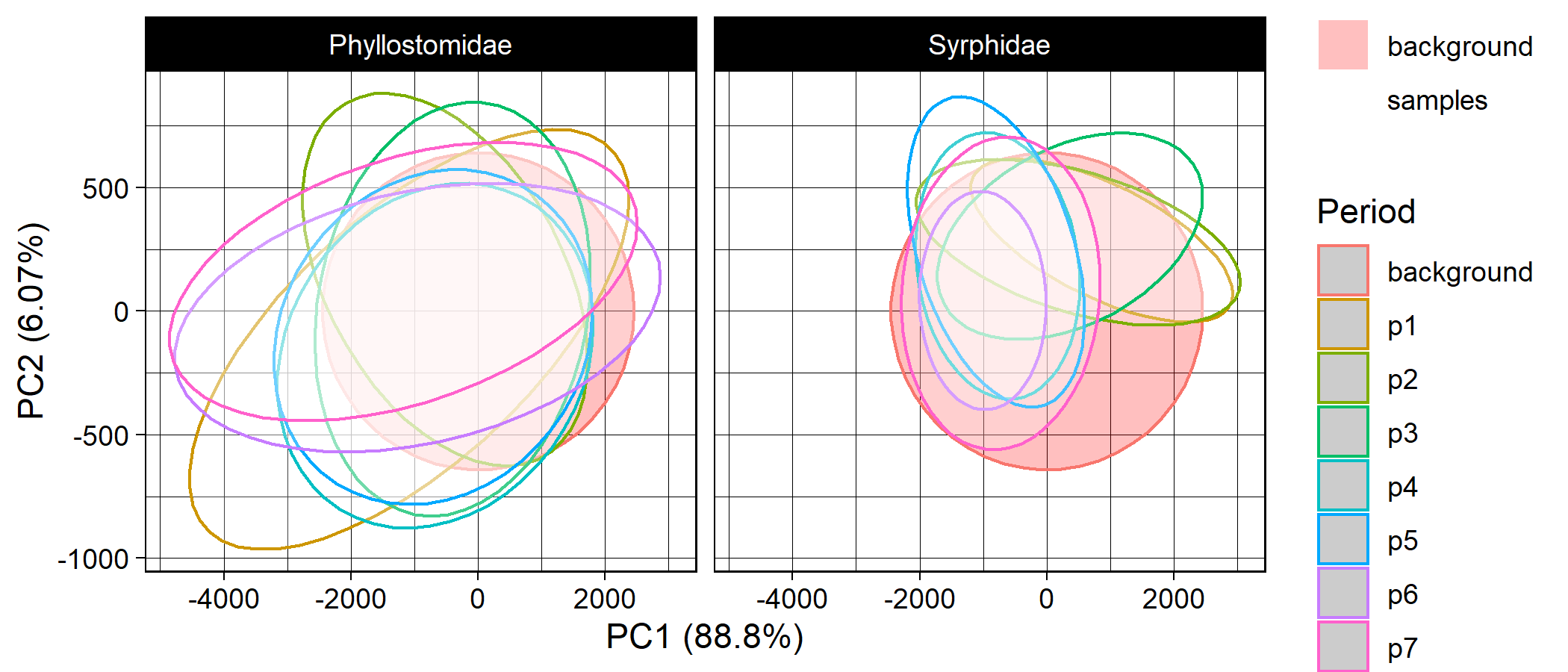

plot of chunk unnamed-chunk-12
