## Supplementary material for "*occAssess*: An R package for assessing potential biases in species occurrence data": vignette 2

Worked example 2: What does a near-random dataset look like?

if (!"occAssess" %in% installed.packages()) devtools::install_github("https://github.com/robboyd/occAssess")
library(occAssess)

### Introduction

This vignette provides a worked example for the functionality of occAssess. For a full tutorial see the main vignette supplied in Boyd et al. (2021).

### Occurrence data

In this worked example, I simulated data that approximates random distributions in space and time for fourty species. The species were simulated with varying prevalence (number of records varies by a factor of 5 with a mean of ~1000). The data were generated at random locations in the UK, and uniformly over the period 2001 to 2010. Each simulated data point was randomly assigned to one of two surveys which are specified in the “identifier” field. The data can be accessed within occAssess as follows:

```

data(random40Species)

spDat <- random40Species

str(spDat)
#> 'data.frame':    40000 obs. of  6 variables:
#>  $ species           : Factor w/ 40 levels "species1","species10",..: 14 19 8 28 5 33 12 27 36 12 ...
#>  $ x                 : num  213500 289500 369500 223500 374500 ...
#>  $ y                 : num  797500 745500 537500 591500 520500 ...
#>  $ year              : int  2009 2004 2006 2010 2009 2004 2004 2004 2010 2001 ...
#>  $ spatialUncertainty: num  9802 3506 15310 9810 7189 ...
#>  $ identifier        : Factor w/ 2 levels "survey1","survey2": 2 2 1 1 1 1 1 1 1 1 ...
```

### Periods

In this example, we will specify five periods over 2001 to 2010

periods <- list(2001:2002, 2003:2004, 2005:2006, 2007:2008, 2009:2010)

### Functions

All of the functions in occAssess require two common arguments: dat and periods (outlined above). I will run through each function in the following, indicating where additional arguments are required. Generally, the functions in occAssess return a list with two elements: one being a ggplot2 object, with a separate panel or feature for each level of identifier; and a second with the data underpinning the plot.

#### assessRecordNumber

The first function I will introduce is the simplest: assessRecordNumber. This function simply plots out the number of records per year in your dataset.

```

nRec <- assessRecordNumber(dat = spDat,
                           periods = periods,
                           species = "species",
                           x = "x",
                           y = "y",
                           year = "year",
                           spatialUncertainty = "spatialUncertainty",
                           identifier = "identifier")

str(nRec$data)
#> 'data.frame':    10 obs. of  3 variables:
#>  $ val   : int  3968 4093 4029 4019 4020 3960 4043 4029 3967 3872
#>  $ group : Factor w/ 2 levels "survey1","survey2": 1 1 1 1 1 2 2 2 2 2
#>  $ Period: int  1 2 3 4 5 1 2 3 4 5

nRec$plot
```

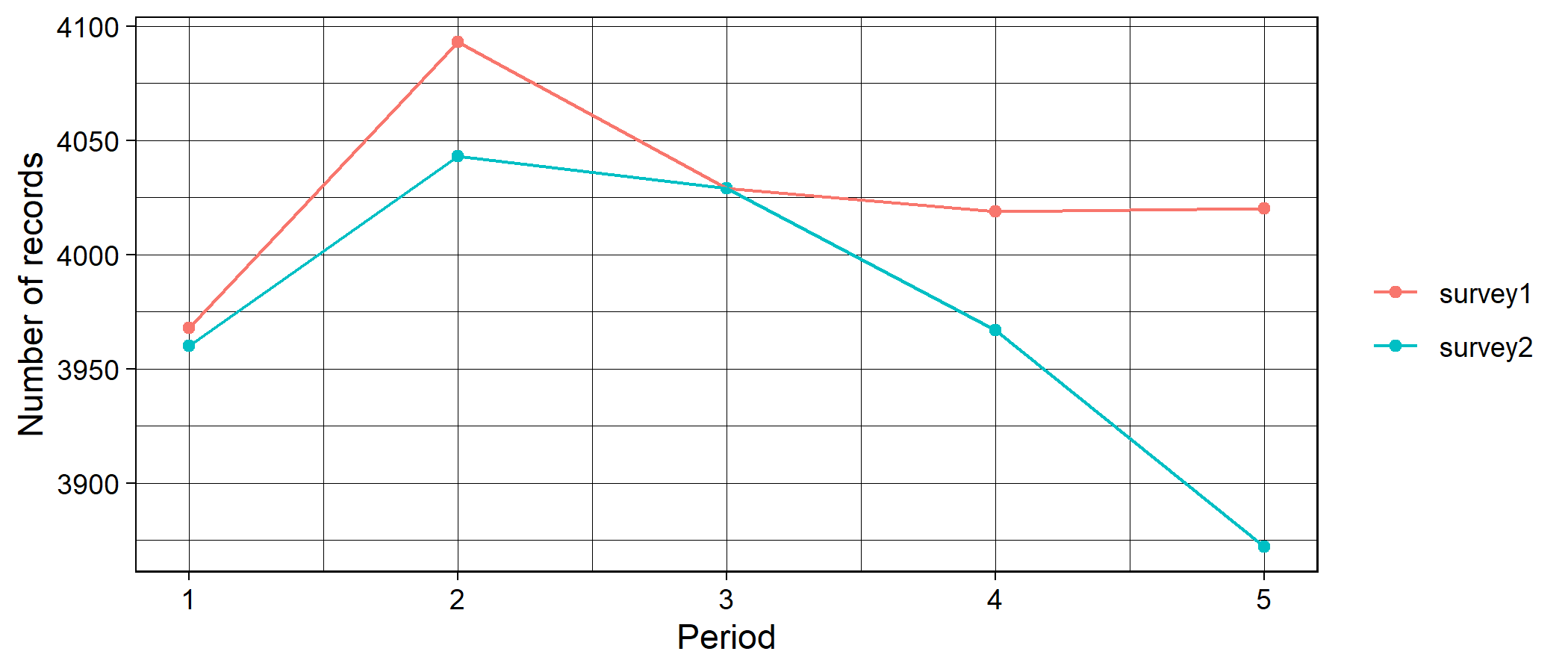

plot of chunk out3

This function enables researchers to quickly establish how the number of records has changed over time.

#### assessSpeciesNumber

In addition to the number of records, you may wish to know how the number of species (taxonomic coverage) in your dataset changes over time. For this you can use the function assessSpeciesNumber:

```

nSpec <- assessSpeciesNumber(dat = spDat,
                           periods = periods,
                           species = "species",
                           x = "x",
                           y = "y",
                           year = "year",
                           spatialUncertainty = "spatialUncertainty",
                           identifier = "identifier")

str(nSpec$data)
#> 'data.frame':    10 obs. of  3 variables:
#>  $ val   : int  40 40 40 40 40 40 40 40 40 40
#>  $ group : Factor w/ 2 levels "survey1","survey2": 1 1 1 1 1 2 2 2 2 2
#>  $ Period: int  1 2 3 4 5 1 2 3 4 5

nSpec$plot
```

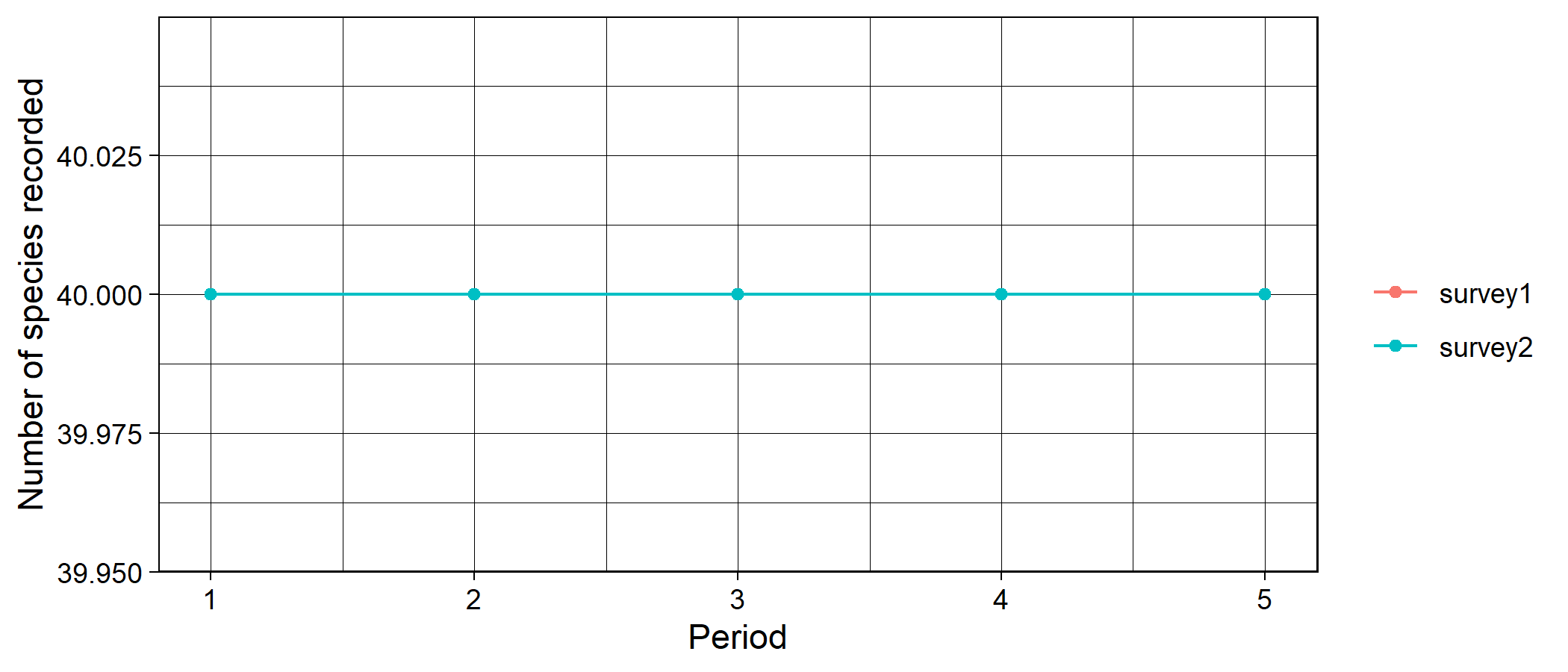

plot of chunk out4

#### assessSpeciesID

It has been speculated that apparent changes in taxonomic coverage could, in fact, reflect a change in taxonomic expertise over time. For example, if fewer individuals have the skill to identify certain species, then it may not appear in your dataset in the later periods. The function assessSpeciesID treats the proportion of species identified to species level as a proxy for taxonomic expertise:

```

propID <- assessSpeciesID(dat = spDat,
                           periods = periods,
                           type = "proportion",
                           species = "species",
                           x = "x",
                           y = "y",
                           year = "year",
                           spatialUncertainty = "spatialUncertainty",
                           identifier = "identifier")

str(propID$data)
#> 'data.frame':    10 obs. of  3 variables:
#>  $ prop  : num  1 1 1 1 1 1 1 1 1 1
#>  $ group : Factor w/ 2 levels "survey1","survey2": 1 1 1 1 1 2 2 2 2 2
#>  $ Period: int  1 2 3 4 5 1 2 3 4 5

propID$plot + ggplot2::ylim(c(0,1)) ## it is easy to modify the outputs of the functions in occAssess to e.g. change axis ranges
```

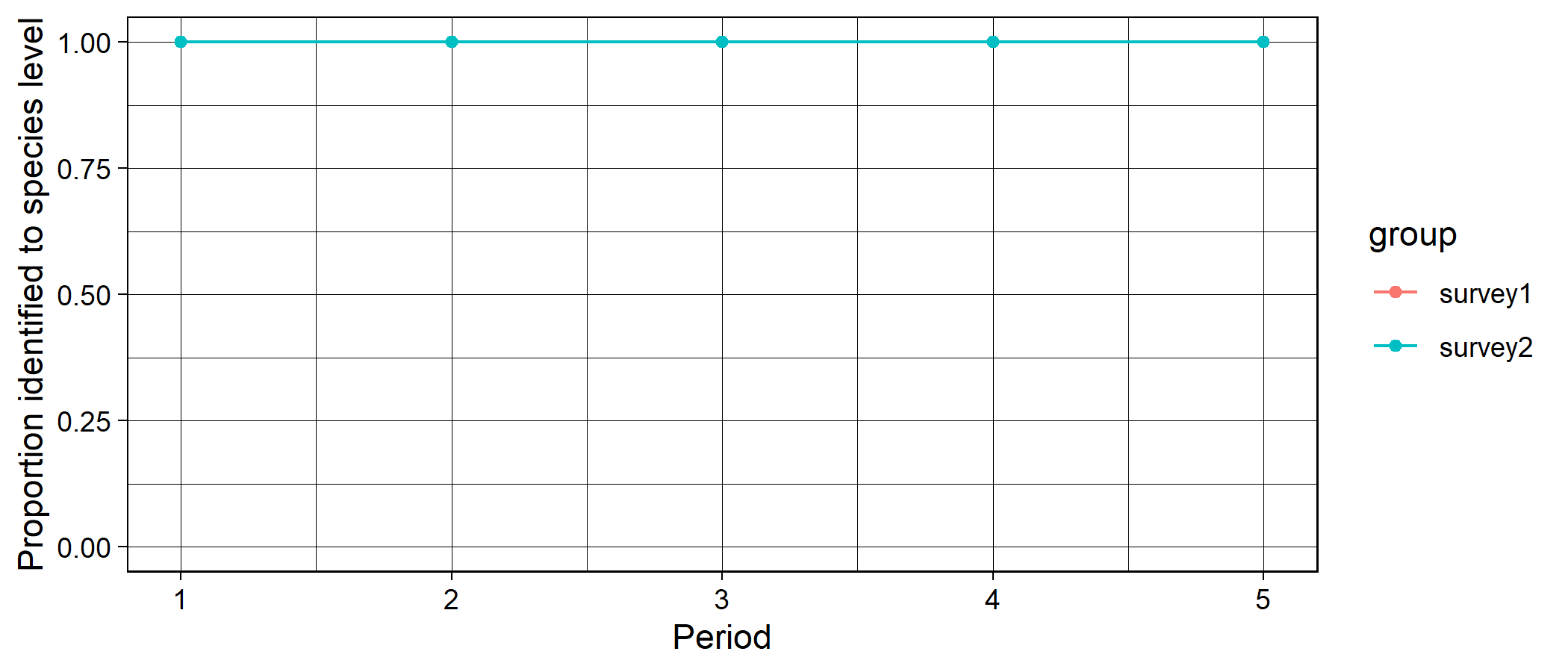

plot of chunk out5

The argument “type” can take the values proportion (proportion of records identified to species level) or count (number of records identified to species level).

#### assessRarityBias

A number of studies have defined taxonomic bias in a dataset as the degree of proportionality between species’ range sizes (usually proxied by the number of grid cells on which it has been recorded) and the total number of records. One can regress the number of records on range size, and the residuals give an index of how over-or undersampled a species is given its prevalence. The function assessRarityBias conducts these analyses for each time period, and uses the r2 value from the linear regressions as an index proportionality between range sizes and number of records. Prevalence may be calculated for each time period if prevPerPeriod = FALSE, and over the whole extent f the data otherwise. Higher values indicate that species’ are sampled in proportion to their range sizes whereas lower values indicate that some species are over- or undersampled.

```

taxBias <- assessRarityBias(dat = spDat,
                            periods = periods,
                            res = 20000,
                            prevPerPeriod = TRUE,
                            species = "species",
                            x = "x",
                            y = "y",
                            year = "year",
                            spatialUncertainty = "spatialUncertainty",
                            identifier = "identifier")

str(taxBias$data)
#> 'data.frame':    10 obs. of  3 variables:
#>  $ period: int  1 2 3 4 5 1 2 3 4 5
#>  $ id    : chr  "survey1" "survey1" "survey1" "survey1" ...
#>  $ index : num  0.994 0.994 0.995 0.995 0.991 ...

taxBias$plot + ggplot2::ylim(c(0,1))
```

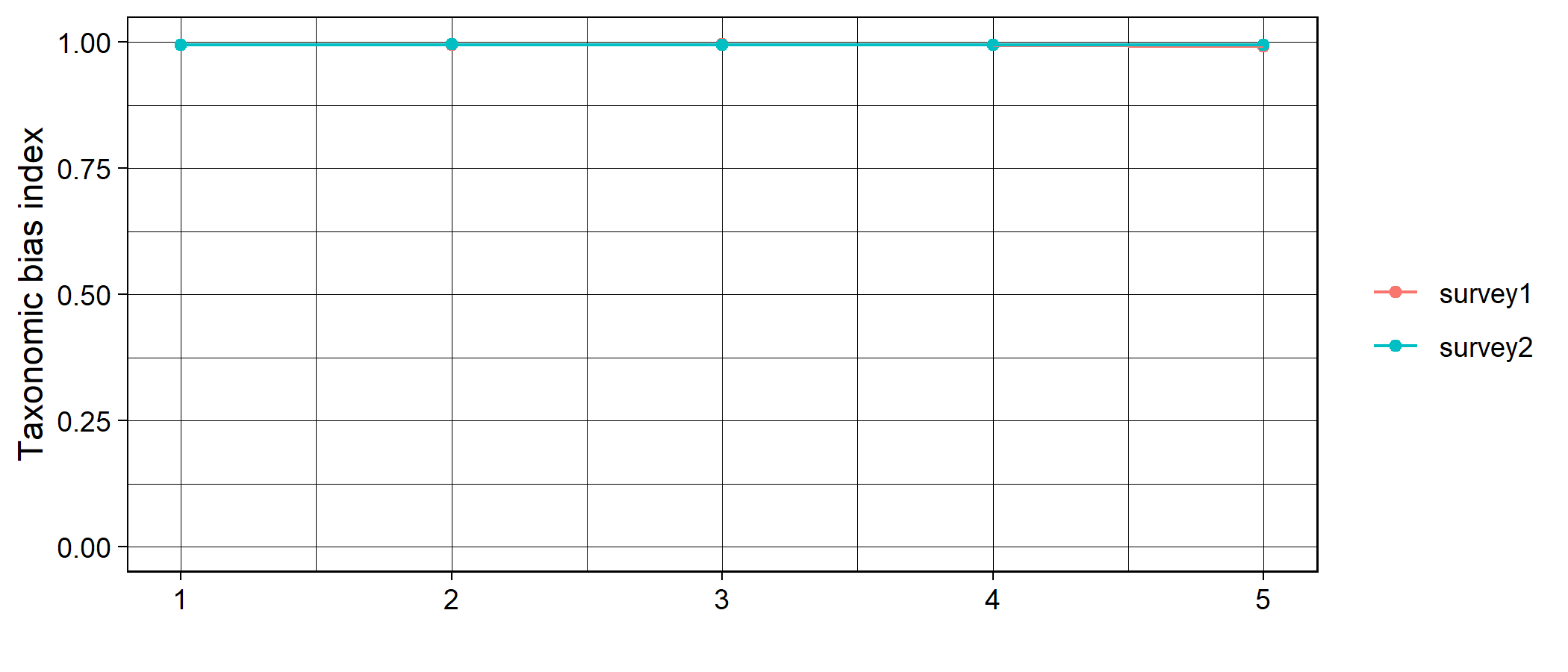

plot of chunk out6

Note the warning message which tells us that there are low numbers of species in some periods (not surprising as the data only contain five species). This represents a small sample size for the regression of range size on number of records so the results should be viewed with caution.

#### assessSpatialCov

The function assessSpatialCov grids your data at a specified spatial resolution then maps it in geographic space. In this example, I provide a shapefile with the boundaries of the UK. If I was working on the WGS84 coordinate reference system (here I am using OSGB 36) this would not be necessary; I could instead use the countries argument and simply specify “UK”. The function returns a list with n elements where n is the number of levels in the identifier field. Each element contains N maps where N is the number of time periods:

```

library(BRCmap) ## a colleague's package that is not publically available. Users will have to provide their own shapefile

data(UK)

shp <- UK[UK$REGION == "Great Britain", ]

head(shp)
#> class       : SpatialPolygonsDataFrame
#> features    : 1
#> extent      : 7422.81, 655616.6, 7041.804, 1218620  (xmin, xmax, ymin, ymax)
#> crs         : +proj=tmerc +lat_0=49 +lon_0=-2 +k=0.9996012717 +x_0=400000 +y_0=-100000 +datum=OSGB36 +units=m +no_defs +ellps=airy +towgs84=446.448,-125.157,542.060,0.1502,0.2470,0.8421,-20.4894
#> variables   : 2
#> names       :        REGION, ID
#> value       : Great Britain,  1

map <- assessSpatialCov(dat = spDat,
                        periods,
                        res = 10000,
                        logCount = TRUE,
                        countries = NULL,
                        shp = shp,
                        species = "species",
                        x = "x",
                        y = "y",
                        year = "year",
                        spatialUncertainty = "spatialUncertainty",
                        identifier = "identifier")
#> Regions defined for each Polygons
#> Regions defined for each Polygons

map$survey1
```

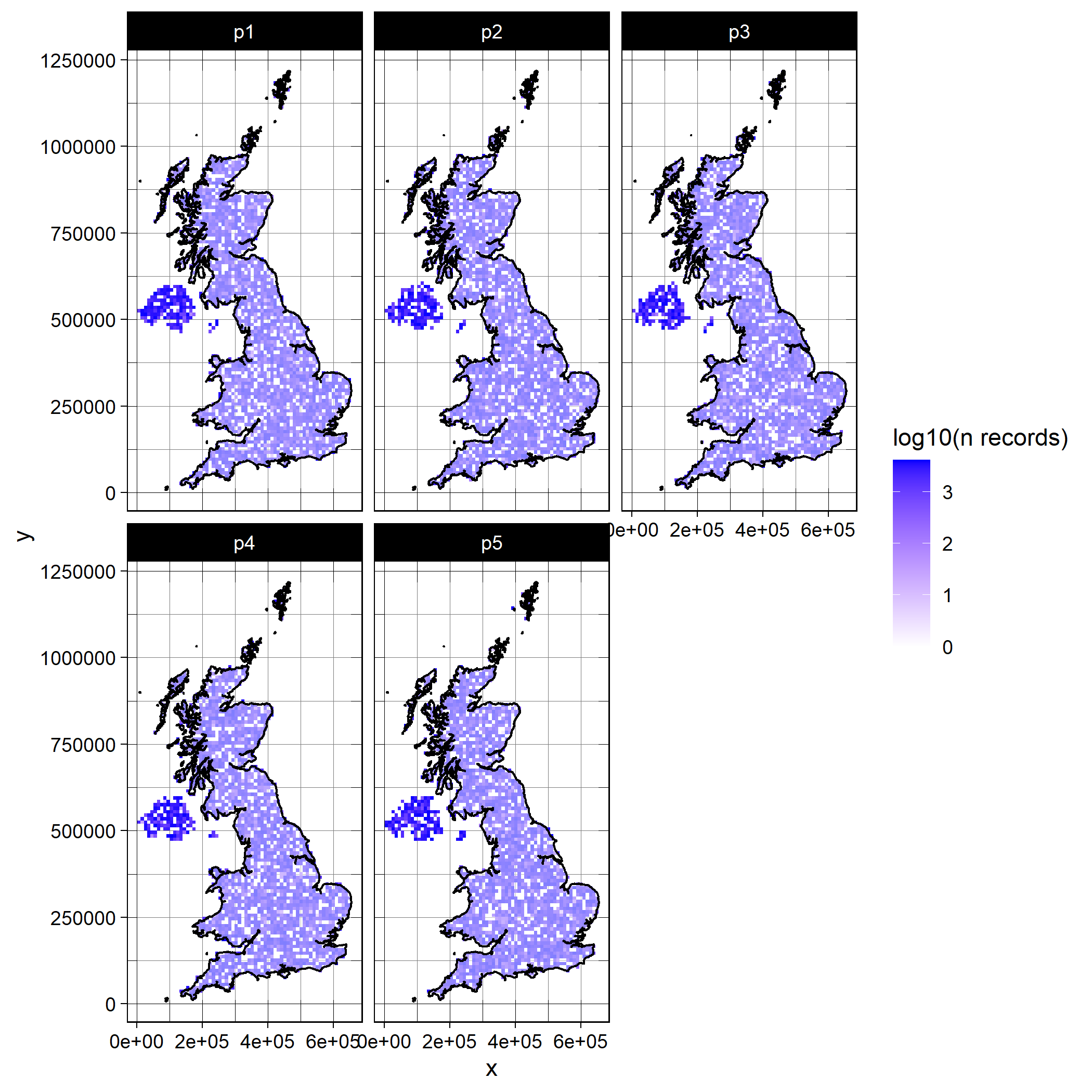

plot of chunk out7

map$survey2

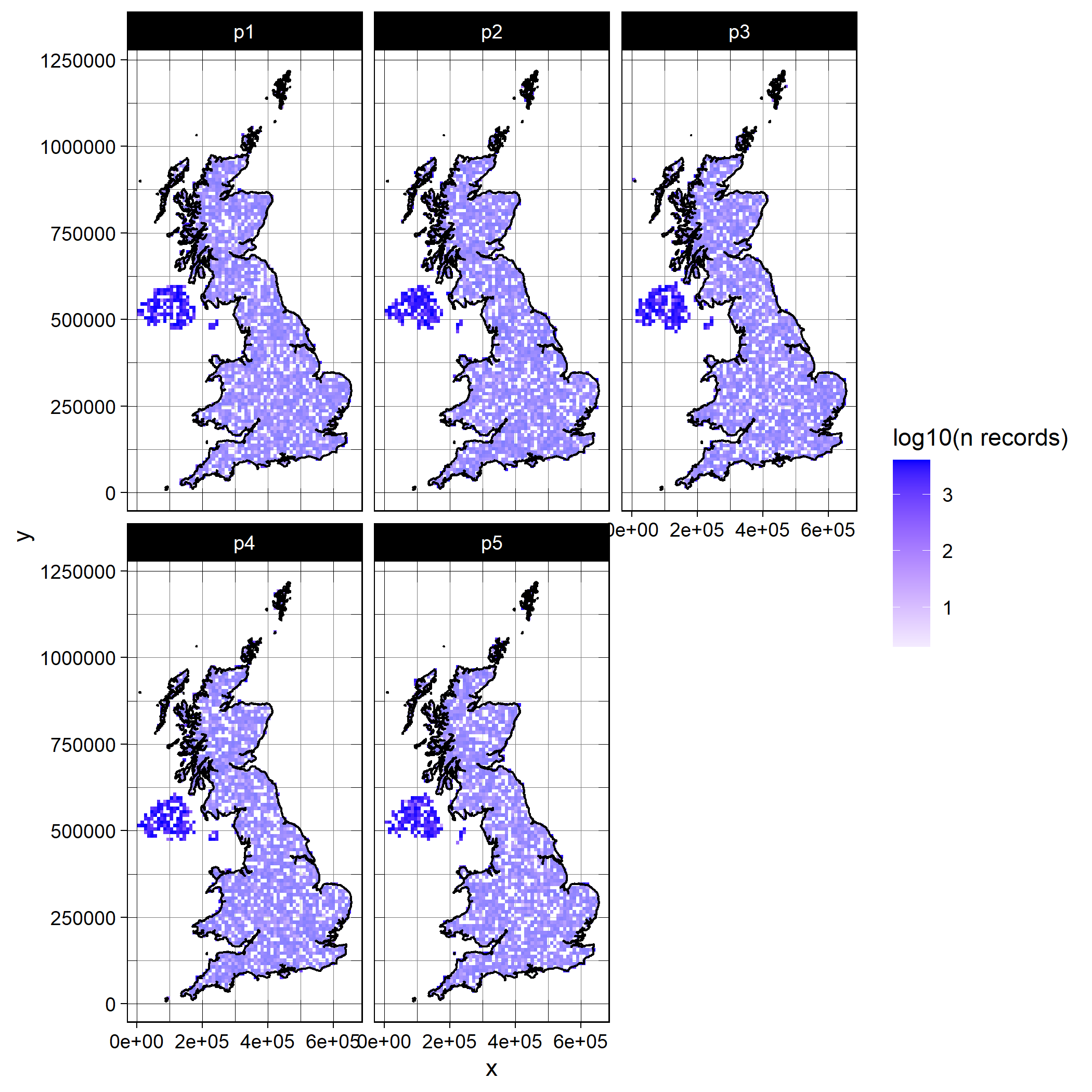

plot of chunk out7

As you can see there are three new arguments to be specified. res is the spatial resolution at which you would like to map the data (units depend on you coordinate reference system, e.g. m if easting and northing, and decimal degress in lon/ lat); logCount indicates whether or not you would like to log10 transform the counts for visual purposes; countries defines the countries covered by your data; and shp is shapefile delimiting your study area if countries are NULL. The countries argument is the simplest way to specify your study boundaries if you are working at country or international level and on the WGS84 lat/ lon coordinate reference system. If you are not working on WGS84, or at a country/ international level, then you will need to provide a shapefile to shp delimiting your boundaries for plotting.

#### assessSpatialBias

Even if your data has good spatial coverage, it may be biased; that is to say, it may deviate from a random distribution in space. The function assessSpatialBias provides an index of how far your data deviates from a random distribution. To do this it simulates an equal number of points to your data randomly across your study region. Then, for each time period, it calculates the average nearest neighbour distance across your data points and divides it by the average nearest neighbour distance from the random sample. If the index is lower than one then your data is more clustered than the random sample, and if it is above one it is more dispersed. To delineate your study area, you must provide a mask layer. The mask is a raster object which is has numeric values within your study area, and is NA outside of your study area. Here, I’ll use some species distribution model outputs for the UK as a mask layer:

```

mask <- raster::raster("C:/Users/Rob.Lenovo-PC/Documents/surpass/Data/Mammals.asc")

mask
#> class      : RasterLayer
#> dimensions : 1250, 700, 875000  (nrow, ncol, ncell)
#> resolution : 1000, 1000  (x, y)
#> extent     : 0, 7e+05, 0, 1250000  (xmin, xmax, ymin, ymax)
#> crs        : NA
#> source     : Mammals.asc
#> names      : Mammals

spatBias <- assessSpatialBias(dat = spDat,
                              periods = periods,
                              mask = mask,
                              nSamps = 10,
                              degrade = TRUE,
                              species = "species",
                              x = "x",
                              y = "y",
                              year = "year",
                              spatialUncertainty = "spatialUncertainty",
                              identifier = "identifier")
#> Warning: `guides(<scale> = FALSE)` is deprecated. Please use `guides(<scale> = "none")` instead.

str(spatBias$data)
#> 'data.frame':    10 obs. of  5 variables:
#>  $ mean      : num  1.009 0.999 1.001 1.001 0.99 ...
#>  $ upper     : num  1.03 1.01 1.02 1.01 1 ...
#>  $ lower     : num  0.991 0.988 0.988 0.987 0.979 ...
#>  $ Period    : chr  "1" "2" "3" "4" ...
#>  $ identifier: chr  "survey1" "survey1" "survey1" "survey1" ...

spatBias$plot + ggplot2::ylim(c(0, 2.15)) # ylim set to theoretical range of possible values for the nearest neighbour index
```

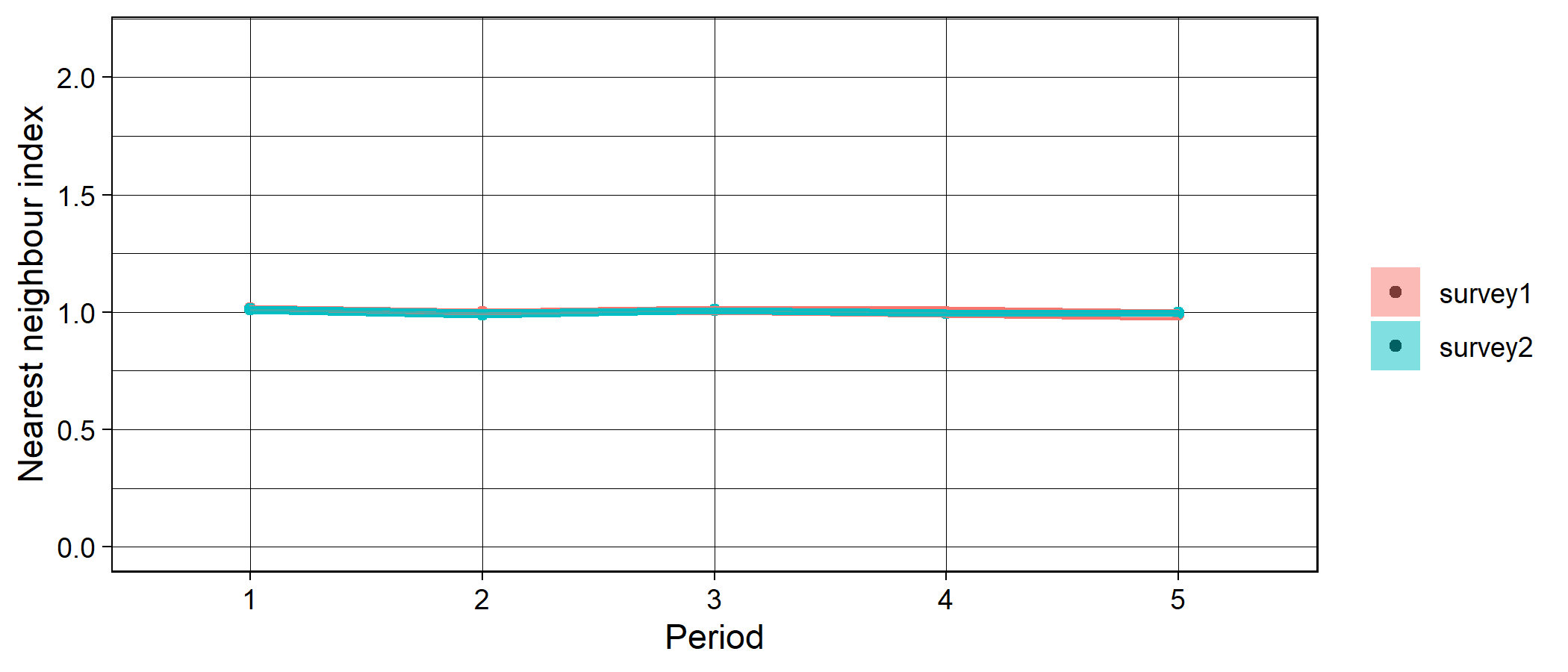

plot of chunk unnamed-chunk-2

The argument nSamps indicates how many random distributions should be drawn, and the argument degrade = TRUE indicates that any duplicated coordinates within a time period and for a given level of identifier are removed. The shaded regions on the plot indicate the 5th and 95th percentiles of the nearest neighbour index calculated over nSamps random samples.

#### assessEnvBias

Spatial bias in your dataset does not necessarily tell you anything about environmental bias. The function assessEnvBias assess the degree to which your data are biased across time periods in environmental space. To do this we first need to get some climate data. I will use the standard suite of 19 bioclimatic variables from worldclim. It is possible to get this data through R using the raster package or online at <https://www.worldclim.org/>. For this example I have extracted the worldclim climate data at the locations of the simulated occurrence data. I have also extracted the climate data at 4000 random locations in the UK as “background” data which I will treat as background climate space. These data can be obtained from within occAssess as follows:

data("random40SpeciesEnvDat") # climate data at locations of simulated occurrence data

data("backgroundEnvDat") # climate data at 4000 random locations

\## How to get the data using raster::getData()

#clim <- raster::getData("worldclim",var="bio",res=10)

\# this will need to be reprojected to the same crs as your data using e.g. raster::projectRaster and then you will need to extract the climate data at the locations of your occurrence data using raster::extract()

assessEnvBias conducts a principal component analysis on your environmental data, then maps your occurrence data in environmental space:

```

envBias <- assessEnvBias(dat = spDat,
                         periods = periods,
                         envDat = random40SpeciesEnvDat,
                         backgroundEnvDat = backgroundEnvDat,
                         xPC = 1,
                         yPC = 2,
                         species = "species",
                         x = "x",
                         y = "y",
                         year = "year",
                         spatialUncertainty = "spatialUncertainty",
                         identifier = "identifier")

envBias$plot
```

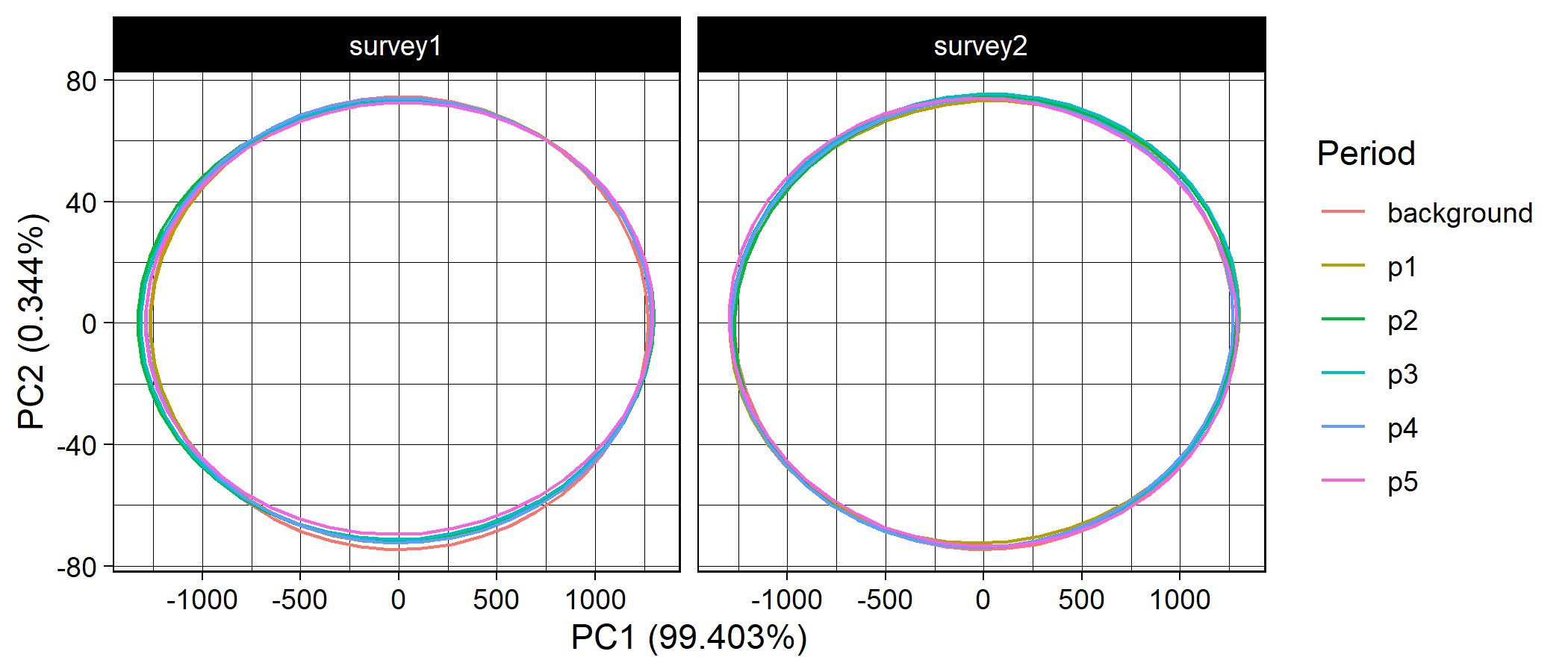
 The arguments yPC and xPC indicate which principal components you would like on the y and x axes, respectively.
