## Supplementary material for "*occAssess*: An R package for assessing potential biases in species occurrence data": vignette 3

Worked example 3: A fully reproducible case study with simulated data

if (!"occAssess" %in% installed.packages()) devtools::install_github("https://github.com/robboyd/occAssess")
library(occAssess)

### Introduction

This vignette provides a worked example for the functionality of occAssess.

### Occurrence data

In this worked example, I simulated data for 11 species across the UK. The species were simulated with varying prevalence (number of records drawn from a uniform distribution between 500 and 1500) and randomly over the period 2001 to 2010. The data were generated as weighted random samples across space (WGS84 coordinate reference system), where the weights were taken from a separate species distribution modelling exercises. Each simulated data point was randomly assigned to one of two surveys which are specified in the “identifier” field. The data can be accessed within occAssess as follows:

```
data("simDat")

spDat <- simDat

str(spDat)
## 'data.frame':    10769 obs. of  6 variables:
##  $ species           : Factor w/ 11 levels "species1","species2",..: 1 1 1 1 1 1 1 1 1 1 ...
##  $ x                 : num  -3.6951 -3.1737 -4.5009 -0.0453 0.8395 ...
##  $ y                 : num  51.2 50.8 58.5 51.5 52.4 ...
##  $ year              : int  2002 2009 2009 2002 2007 2010 2010 2008 2002 2010 ...
##  $ spatialUncertainty: num  19216 19239 14255 11704 7791 ...
##  $ identifier        : Factor w/ 2 levels "survey1","survey2": 2 1 1 1 2 2 2 1 1 1 ...
```

### Periods

In this example, we will specify five periods over 2001 to 2010

periods <- list(2001:2002, 2003:2004, 2005:2006, 2007:2008, 2009:2010)

### Functions

All of the functions in occAssess require two common arguments: dat and periods (outlined above). I will run through each function in the following, indicating where additional arguments are required. Generally, the functions in occAssess return a list with two elements: one being a ggplot2 object, with a separate panel or feature for each level of identifier; and a second with the data underpinning the plot.

#### assessRecordNumber

The first function I will introduce is the simplest: assessRecordNumber. This function simply plots out the number of records per period in your dataset. Note that, for all functions, users must specify six arguments corresponding to the six mandatory fields in occAssess: species, x, y, year, spatialuncertainty and identifier. These arguments indicate which columns in dat apply to each field.

```
nRec <- assessRecordNumber(dat = spDat,
                           periods = periods,
                           species = "species",
                           x = "x",
                           y = "y",
                           year = "year",
                           spatialUncertainty = "spatialUncertainty",
                           identifier = "identifier")


nRec$plot
```

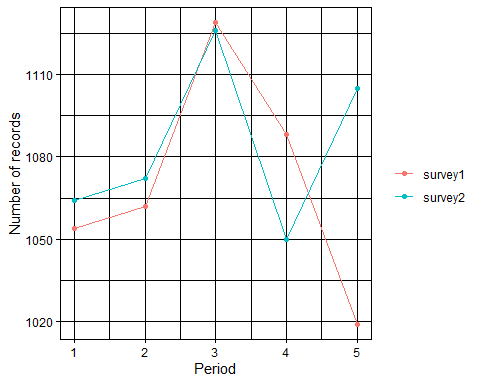

This function enables researchers to quickly establish how the number of records has changed over time.

#### assessSpeciesNumber

In addition to the number of records, you may wish to know how the number of species (taxonomic coverage) in your dataset changes over time. For this you can use the function assessSpeciesNumber:

```
nSpec <- assessSpeciesNumber(dat = spDat,
                           periods = periods,
                           species = "species",
                           x = "x",
                           y = "y",
                           year = "year",
                           spatialUncertainty = "spatialUncertainty",
                           identifier = "identifier")

nSpec$plot
```

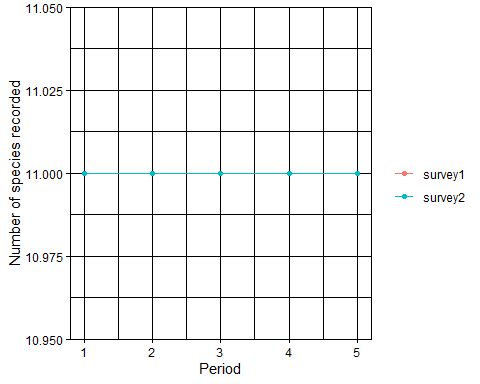

#### assessSpeciesID

It has been speculated that apparent changes in taxonomic coverage could, in fact, reflect a change in taxonomic expertise over time. For example, if fewer individuals have the skill to identify certain species, then it may not appear in your dataset in the later periods. The function assessSpeciesID treats the proportion of records identified to species level as a proxy for taxonomic expertise:

```
propID <- assessSpeciesID(dat = spDat,
                           periods = periods,
                           type = "proportion",
                           species = "species",
                           x = "x",
                           y = "y",
                           year = "year",
                           spatialUncertainty = "spatialUncertainty",
                           identifier = "identifier")

propID$plot + ggplot2::ylim(c(0,1)) ## it is easy to modify the outputs of the functions in occAssess to e.g. change axis ranges
```

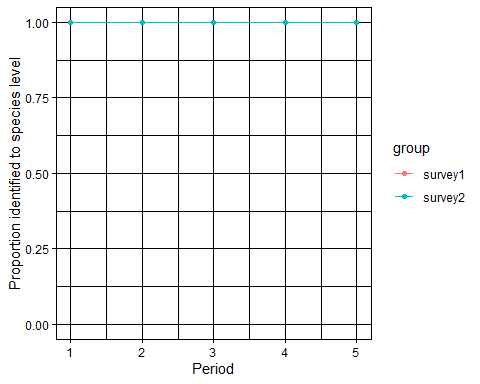

The argument “type” can take the values proportion (proportion of records identified to species level) or count (number of records identified to species level).

#### assessRarityBias

A number of studies have defined taxonomic bias in a dataset as the degree of proportionality between species’ range sizes (usually proxied by the number of grid cells on which it has been recorded) and the total number of records. One can regress the number of records on range size, and the residuals give an index of how over-or undersampled a species is given its prevalence. The function assessRarityBias conducts these analyses for each time period, and uses the r2 value from the linear regressions as an index proportionality between range sizes and number of records. Prevalence may be calculated for each time period if prevPerPeriod = FALSE, and over the whole extent f the data otherwise. Higher values indicate that species’ are sampled in proportion to their range sizes whereas lower values indicate that some species are over- or undersampled.

```
taxBias <- assessRarityBias(dat = spDat,
                            periods = periods,
                            res = 0.5,
                            prevPerPeriod = FALSE,
                            species = "species",
                            x = "x",
                            y = "y",
                            year = "year",
                            spatialUncertainty = "spatialUncertainty",
                            identifier = "identifier")

taxBias$data
##    period      id        index
## 1       1 survey1 5.439108e-03
## 2       2 survey1 3.819923e-03
## 3       3 survey1 1.013494e-02
## 4       4 survey1 1.277338e-03
## 5       5 survey1 9.356627e-03
## 6       1 survey2 6.180651e-02
## 7       2 survey2 1.791968e-04
## 8       3 survey2 1.941553e-05
## 9       4 survey2 5.543759e-03
## 10      5 survey2 1.639747e-02
taxBias$plot + ggplot2::ylim(c(0,1))
```

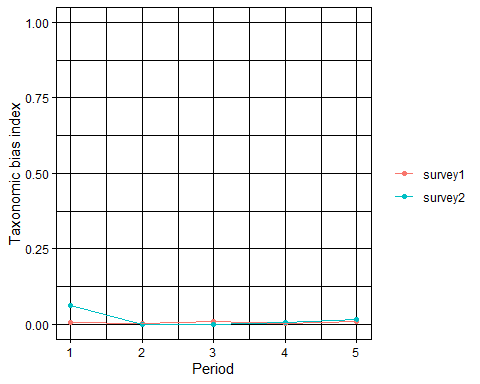

As you can see, the rarity bias index is roughly 0 for both levels of identifier and in each period. This indicates that species are not recorded in proportion to their commonness, which is expected given that the data were not simulated in such a way.

#### assessSpatialCov

The function assessSpatialCov grids your data at a specified spatial resolution then maps it in geographic space. As I am working on the WGS84 coordinate reference system, I do not have to provide a shapefile with the relevant country borders; instead, I can specify “UK” in the countries argument. The function returns a list with n elements where n is the number of levels in the identifier field. Each element contains N maps where N is the number of time periods:

```
map <- assessSpatialCov(dat = spDat,
                        periods = periods,
                        res = 0.5,
                        logCount = TRUE,
                        countries = "UK",
                        species = "species",
                        x = "x",
                        y = "y",
                        year = "year",
                        spatialUncertainty = "spatialUncertainty",
                        identifier = "identifier")

map$survey1
```

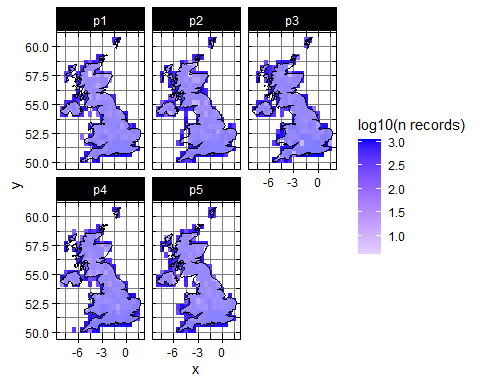

map$survey2

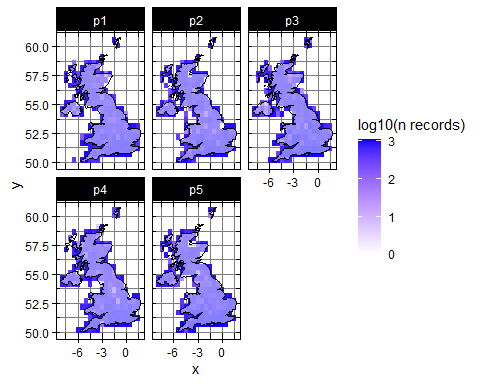

As you can see there are three new arguments to be specified. res is the spatial resolution at which you would like to map the data (units depend on you coordinate reference system, e.g. m if easting and northing, and decimal degress in lon/ lat); logCount indicates whether or not you would like to log10 transform the counts for visual purposes; countries defines the countries covered by your data; and shp is shapefile delimiting your study area if countries are NULL. The countries argument is the simplest way to specify your study boundaries if you are working at country or international level and on the WGS84 lat/ lon coordinate reference system. If you are not working on WGS84, or at a country/ international level, then you will need to provide a shapefile to shp delimiting your boundaries for plotting.

#### assessSpatialBias

Even if your data has good spatial coverage, it may be biased; that is to say, it may deviate from a random distribution in space. The function assessSpatialBias provides an index of how far your data deviates from a random distribution. To do this it simulates an equal number of points to your data randomly across your study region. Then, for each time period, it calculates the average nearest neighbour distance across your data points and divides it by the average nearest neighbour distance from the random sample. If the index is lower than one then your data is more clustered than the random sample, and if it is above one it is more dispersed. To delineate your study area, you must provide a mask layer. The mask is a raster object which is has numeric values within your study area, and is NA outside of your study area. You can access a mask layer from within occAssesss for this example:

```
data("UKWGS84mask")

UKWGS84mask
## class      : RasterLayer
## dimensions : 1284, 829, 1064436  (nrow, ncol, ncell)
## resolution : 0.0158, 0.00898  (x, y)
## extent     : -9.469967, 3.628233, 49.68321, 61.21353  (xmin, xmax, ymin, ymax)
## crs        : +proj=longlat +datum=WGS84 +no_defs
## source     : memory
## names      : layer
## values     : 1, 1  (min, max)
spatBias <- assessSpatialBias(dat = spDat,
                              periods = periods,
                              mask = UKWGS84mask,
                              nSamps = 10,
                              degrade = TRUE,
                              species = "species",
                              x = "x",
                              y = "y",
                              year = "year",
                              spatialUncertainty = "spatialUncertainty",
                              identifier = "identifier")
## Warning: `guides(<scale> = FALSE)` is deprecated. Please use `guides(<scale> =
## "none")` instead.
str(spatBias$data)
## 'data.frame':    10 obs. of  5 variables:
##  $ mean      : num  0.96 0.953 0.925 0.956 0.99 ...
##  $ upper     : num  0.997 0.97 0.952 0.973 1.016 ...
##  $ lower     : num  0.928 0.933 0.908 0.93 0.969 ...
##  $ Period    : chr  "1" "2" "3" "4" ...
##  $ identifier: chr  "survey1" "survey1" "survey1" "survey1" ...
spatBias$plot + ggplot2::ylim(c(0, 2.15)) # ylim set to theoretical range of possible values for the nearest neighbour index
```

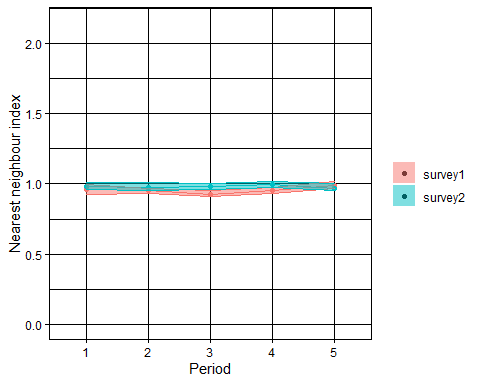

The argument nSamps indicates how many random distributions should be drawn, and the argument degrade = TRUE indicates that any duplicated coordinates within a time period and for a given level of identifier are removed. The shaded regions on the plot indicate the 5th and 95th percentiles of the nearest neighbour index calculated over nSamps random samples.

#### assessEnvBias

Spatial bias in your dataset does not necessarily tell you anything about environmental bias. The function assessEnvBias assess the degree to which your data are biased across time periods in environmental space. To do this we first need to get some climate data. I will use the standard suite of 19 bioclimatic variables from worldclim. It is possible to get this data through R using the raster package or online at <https://www.worldclim.org/>. For this example, you can access the environmental data that corresponds to the occurrence data from within occAssess. I have also extracted the climate data at 4000 random locations in the UK as “background” data which I will treat as background climate space. These data can be obtained as follows:

data("simEnvDat") # climate data at locations of simulated occurrence data

data("backgroundEnvDat") # climate data at 4000 random locations across the UK

\## How to get the data using raster::getData()

#clim <- raster::getData("worldclim",var="bio",res=10)

\# this will need to be reprojected to the same crs as your data using e.g. raster::projectRaster and then you will need to extract the climate data at the locations of your occurrence data using raster::extract()

assessEnvBias conducts a principal component analysis on your environmental data, then maps your occurrence data in environmental space:

```
envBias <- assessEnvBias(dat = spDat,
                         periods = periods,
                         envDat = simEnvDat,
                         backgroundEnvDat = backgroundEnvDat,
                         xPC = 1,
                         yPC = 2,
                         species = "species",
                         x = "x",
                         y = "y",
                         year = "year",
                         spatialUncertainty = "spatialUncertainty",
                         identifier = "identifier")

envBias$plot
```

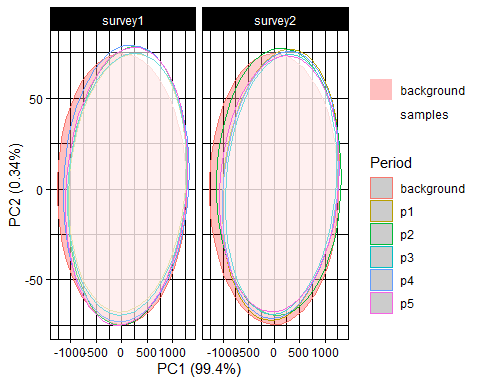

The arguments yPC and xPC indicate which principal components you would like on the y and x axes, respectively.
